# Single cell landscape of hypertrophic scars identifies serine proteases as key regulators of myofibroblast differentiation

**DOI:** 10.1101/2020.06.17.157073

**Authors:** Vera Vorstandlechner, Maria Laggner, Dragan Copic, Yiyan Chen, Bahar Golabi, Werner Haslik, Christine Radtke, Erwin Tschachler, Hendrik Jan Ankersmit, Michael Mildner

**Affiliations:** Laboratory for Cardiac and Thoracic Diagnosis, Regeneration and Applied Immunology, Division of Thoracic Surgery, Medical University of Vienna; Aposcience AG, Vienna, Austria; Division of Plastic and Reconstructive Surgery, Department of Surgery, Medical University of Vienna, Vienna, Austria; Department of Dermatology, Medical University of Vienna, Vienna, Austria

## Abstract

Despite recent advances in understanding skin scarring, mechanisms triggering hypertrophic scar formation are still poorly understood. In the present study we performed single-cell sequencing of mature human hypertrophic scars and developing scars in mice.

Compared to normal skin, we found significant differences in gene expression in most cell types present in scar tissue. Fibroblasts (FBs) showed the most prominent alterations in gene expression, displaying a distinct fibrotic signature. By comparing genes upregulated in murine FBs during scar development with genes highly expressed in mature human hypertrophic scars, we identified a group of serine proteases, tentatively involved in scar formation. Two of them, dipeptidyl-peptidase 4 (DPP4) and urokinase (PLAU), were further analyzed in functional assays, revealing a role in TGFβ1-mediated myofibroblast differentiation and over-production of components of the extracellular matrix (ECM) without interfering with the canonical TGFβ1-signaling pathway.

In this study, we delineate the genetic landscape of hypertrophic scars and present new insights into mechanisms involved in hypertrophic scar formation. Our data suggest the use of serine protease inhibitors for the treatment of skin fibrosis.

## Introduction

Hypertrophic scars are a complex medical problem and a significant global disease burden (Bayat et al, 2003; Leavitt et al, 2016). In the western world, an estimated number of 100 million people develop scars every year, approximately 11 million of which bear keloid scars and 4 million suffer from burn scars (Bayat et al, 2003). In the USA, an estimated amount of 12 Billion Dollars is spent annually on the treatment of skin scarring (Sen et al, 2009). For affected persons, a pathological hypertrophic scar causes significant functional impairment, pain, pruritus, and a reduction in quality of life (Bock et al, 2006; Van Loey et al, 2008).

Wound healing is a tightly coordinated, three-step process, characterized by an acute inflammatory phase, a proliferative phase, and a remodeling phase. Prolonged inflammation results in increased fibroblast (FB) activity, with enhanced secretion of transforming growth factor beta 1 (TGFβ1), TGFβ 2, insulin-like growth factor (IGF1), and other cytokines (Lee & Jang, 2018). TGFβ1 drives differentiation of FBs into myofibroblasts, which have a contractile phenotype, are characterized by excessive secretion of ECM-components (Hinz, 2016), and are the major contributors to the formation of hypertrophic scars (Nabai et al, 2020). Mature hypertrophic scars display strong tissue contraction (Lee & Jang, 2018), and dense, parallel or whorl-like ECM (Nabai et al, 2020).

Topical silicone application, compression or massage therapy, laser ablation, surgery, and intralesional injection of triamcinolone (TAC), corticosteroids, or 5-Fluorouracil (5-FU) are the most commonly used options for prevention or treatment of hypertrophic scars (Anthonissen et al, 2016; Kafka et al, 2017; Lee & Jang, 2018; Tredget et al, 2017). However, many of these therapies lack evidence of efficacy and safety, and mechanisms of actions are still unclear (Bao et al, 2019; Kafka et al, 2017).

Recently, several proteases became the focus of drug development in fibrotic diseases, as they have been shown to be involved in ECM-breakdown and the activation of growth factors in tissue remodeling (Kanno, 2019; Menou et al, 2018). Serine proteases/peptidases constitute a large, diverse group of proteases, divided into 13 clans and 40 families (Page & Di Cera, 2008).The group of trypsins comprises proteases contributing to vital processes such as blood coagulation, fibrinolysis, apoptosis and immunity (Di Cera, 2009). Members of this family include urokinase, HTRA1/3 (high temperature requirement A1/3 peptidase), several coagulation factors and complement components, PRSS-like serine proteases, granzymes, and cathepsin G (Di Cera, 2009; Rawlings & Barrett, 1999). Inhibitors of PLAU have been shown to counteract fibrotic processes in cardiac and pulmonary fibrosis in human *in vitro* studies and in mouse experiments (Gupta & Donahue, 2017; Schuliga et al, 2017). Recently, the serine protease DPP4 became the center of attention, since DPP4 inhibitors (gliptins) have been clinically used for the treatment of diabetes mellitus (Makrilakis, 2019). DPP4 was also implicated in a variety of fibrotic pathologies, including cardiac, hepatic, renal, and dermal fibrosis (Aroor et al, 2017; Hong et al, 2017; Kaji et al, 2014; Suzuki et al, 2017; Uchida et al, 2017) and inhibition of DPP4 activity mitigated fibrotic processes in animal models (Gupta & Donahue, 2017; Hu & Longaker, 2016; Lay et al, 2019; Schuliga et al, 2018; Schuliga et al, 2017; Shi et al, 2016). However, the contribution to human scar formation and the underlying anti-fibrotic mechanisms are so far not known.

Here, we used the new approach of scRNAseq to thoroughly study gene expression and mechanisms involved in hypertrophic scar formation. We aimed to identify new genes regulated in scar tissue, and to uncover potential new targets for drug development towards scar-free wound healing or full reversion of a present scar.

## Methods

### Ethical statement

The Vienna Medical University ethics committee approved the use of healthy abdominal skin (Vote Nr. 217/2010) and of scar tissue (Vote Nr. 1533/2017) and all donors provided written informed consent. Animal experiments were approved by the Medical University of Vienna ethics committee and by the Austrian Federal Ministry of Education, Science and Research (Vote Nr. BMBWF-66.009/0075-V/3b/2018) and performed in accordance with the Austrian guidelines for the use and care of laboratory animals.

### Scar and skin samples

Resected scar tissue was obtained from patients who underwent elective scar resection surgery (donor information is provided in Table S1). Scars were classified as hypertrophic, pathological scars according to POSAS (Fearmonti et al, 2011) by a plastic surgeon. Only mature scars, which had not been treated before and persisted for more than two years were used for all experiments. All donors had no known chronic diseases and received no chronic medication. Healthy skin was obtained from female donors between 25-45 years from surplus abdominal skin removed during elective abdominoplasty.

### Mouse full skin wounding and scar maturation

Female Balb/c mice bred at the animal facility of the Medical University of Vienna (Himberg, Austria) were housed under specific-pathogen-free conditions with 12h/12h light/dark cycles and food and water access *ad libidum.* For full thickness skin wounds, mice were anesthetized with 100 mg/kg Xylazin and 5 mg/kg ketamin (both Sigma-Aldrich, St. Louis, MO, USA) intraperitoneally. Postoperative analgesia was provided with 0.1 mg/kg Buprenorphin (Temgesic^®^, Indivior Inc., North Chesterfield, VA, USA) subcutaneously and 0.125 mg/ml Piritramid (Janssen-Cilag Pharma, Vienna, Austria) in drinking water *ad libidum.* A 9×9 mm^2^ area was marked on shaved backs and excised with sharp scissors. The wounds were left to heal uncovered without any further intervention. Mice were sacrificed 6 or 8 weeks after wounding, and scar tissues were isolated.

### Single cell isolation and fluorescence-activated cell sorting (FACS)

Four mm biopsies were taken from human skin, human scar, and from mouse scar tissue, and enzymatically digested with MACS Miltenyi Whole Skin Dissociation Kit (Miltenyi Biotec, Bergisch-Gladbach, Germany) for 2.5h according to the manufacturer’s protocol. After processing on a GentleMACS OctoDissociator (Miltenyi), cell suspensions were passed through a 70 μm and a 40 μm filter and stained with DAPI nuclear dye. Cells were sorted on a MoFlo Astrios high speed cell sorting device (Beckman-Coulter, Brea, CA, USA), and only DAPI-negative cells, representing viable cells, were used for single cell RNAseq.

### Generation of single-cell gel-bead in emulsions (GEMs) and library preparation

Immediately after sorting, viable cells were loaded on a 10X-chromium instrument (Single cell gene expression 3’v2/3, 10X Genomics, Pleasanton, CA, USA) to generate GEMs. GEM-generation, library preparation, RNA-sequencing, demultiplexing and counting were done by the Biomedical Sequencing Core Facility of the Center for Molecular Medicine (CeMM, Vienna, Austria). Sequencing was performed on an Illumina HiSeq 3000/4000 (Illumina, San Diego, CA, USA) with 3 samples per lane, 2×75bp, and paired-end sequencing.

### Cell-gene matrix preparation and downstream analysis

Raw sequencing files were demultiplexed, aligned to the human or mouse reference genome (GrCh38/ mm10) and counted using the Cellranger pipelines (Cellranger v3, 10X Genomics). The resulting cell-gene matrices were processed using the ‘Seurat’-package (Seurat v3.1.0, Satija Lab, New York, NY, USA) in R-studio in R (R v3.6.2, The R Foundation, Vienna, Austria). From each sample, unwanted variations and low-quality cells were filtered by removing cells with high and low (>3000 and <200) Unique Molecular Identifier (UMI)-counts. First, healthy skin and scar samples were integrated separately to avoid clustering according to donors, and for batch correction. Subsequently, skin and scar data were integrated again into one dataset. Data integration was performed according to the recommended workflow by Butler *et al.* and Stuart *et al.* (Butler et al, 2018; Stuart et al, 2018). After quality control comparing all donors, we obtained transcriptome data from a total of 25.083 human skin and scar cells, with a median of 24943 reads and 851 detected genes per cell. In mice, we obtained data from 6561 cells 6 weeks after wounding, and 9393 cells 8 weeks after wounding. The samples displayed a median of 24774 reads per cell, and median of 1969 detected genes per cell. After quality control, all mouse samples were integrated together in one integration step. In both datasets, normalized count numbers were used for differential gene expression analysis, for visualization in violin plots, feature plots, dotplots and heatmaps, when displaying features that vary across conditions, as recommended by current guidelines (Luecken & Theis, 2019). In both datasets, cell types were identified by well-established marker gene expression (Figure S1A, Figure S4A). For identification of differentially expressed genes (DEGs), normalized count numbers were used, including genes present in the integrated dataset to avoid calculation of batch effects. As keratin and collagen genes were previously found to contaminate skin biopsy datasets and potentially provide a false-positive signal (Rojahn et al, 2020), these genes *(COL1A1, COL1A2, COL3A1* and *KRT1 KRT5, KRT10, KRT14, KRTDAP)* were excluded from DEG calculation in non-fibroblast clusters (collagens) or non-keratinocyte clusters (keratins), respectively. Moreover, genes *Gm42418, Gm17056* and *Gm26917* caused technical background noise and batch effect in mouse scRNAseq, as described before (Hammond et al, 2019), and were thus excluded from the dataset.

### Pseudotime analyses

Pseudotime analyses, trajectory-construction and calculation of pseudotime-dependent gene expression were performed in Monocle 2 (Monocle2, v2.14.0, Trapnell Lab, University of Washington, Seatlle, WA, USA) (Qiu et al, 2017a; Trapnell et al, 2014). From the integrated FB subset Seurat-object, data were converted into a monocle-compatible CellDataSet. Analysis was then performed according to the recommended pipeline. Cells with mRNA counts two standard deviations above or below the mean were excluded. Size factors and dispersions were estimated, tSNE-reduction and clustering were performed (Qiu et al, 2017a; Qiu et al, 2017b; Trapnell et al, 2014). As input for pseudotime ordering, differentially expressed genes between skin and scar were used, and trajectories were constructed with DDRTree (R-package ‘DDRTree’ v0.1.5, by Qi Mao, Li Wang *et al.*, 2015) (Qiu et al, 2017b).

### STRING functional protein interaction networks

Protein interaction networks were generated by imputing a list of genes (e.g. upregulated genes in FBs scar vs skin) into the String v11.0 online tool (Snel et al, 2000; Szklarczyk et al, 2019) (https://string-db.org/, String consortium, Swiss Institute of Bioinformatics/EMBL Heidelberg/University of Zurich/Novo Nordisk Foundation CPR).

### Gene ontology (GO)-networks

Gene lists of significantly regulated genes (adjusted p-value <0.05, average log fold change [avg_logFC] >0.1) were imputed in ClueGO v2.5.5 (Bindea et al, 2009) plug-in in Cytoscape v3.7.2 (Lotia et al, 2013) with medium GO-specificity, with GO-term fusion, and only significant *(P* value <.05) GO-terms are depicted as circles, whereby circle size correlates with *P* value, and lines represent functional connection of respective GO-terms.

### Immunfluorescence stainings

Immmunofluorescence staining on formalin-fixed, paraffin-embedded (FFPE) sections of skin and scar tissue were performed as described previously (Gschwandtner et al, 2013). Antibodies were used as indicated in Table S2.

### RNAScope in situ hybridization

FFPE-sections of human skin and scar tissue were prepared according to RNAScope (ACDBio, Bio-Techne, Bristol, UK) pre-treatment protocol, hybridized with probes targeting human DPP4 (RNAscope^®^ Probe-Hs-DPP4) and PLAU (RNAscope^®^ Probe-Hs-PLAU), and visualized with RNAscope 2.5 HD Assay – RED as suggested by the manufacturer. Images were acquired by AX70 microscope (Olympus, Tokyo, Japan) using the imaging software MetaMorph (Olympus).

### Isolation of primary skin fibroblasts

Five mm biopsies were taken from fresh abdominal skin, washed in phosphate-buffered saline (PBS), and incubated in 2.4 U/ml Dispase II (Roche, Basel, Switzerland) overnight at 4°C. The next day, epidermis was separated from dermis, and dermis was incubated with Liberase TM (Merck Millipore, Burlington, MA, USA) in Dulbeccos modified eagle medium (DMEM, Thermo Fisher Scientific, Waltham, MA, USA) without supplements at 37°C for 2h. Next, the dermis was passed through 100 μm and 40 μm filters, rinsed with PBS, and cells were plated in a T175 cell culture flask. Medium was changed the next day, and then every other day until FBs reached 90% confluency. First passage FBs were used for TGFβ1-stimulation experiments.

### TGFβ 1-induced myofibroblast differentiation

After the first passage, isolated primary FBs were plated in 6-well plates, supplied with DMEM + 10% fetal bovine serum (FBS, Thermo Fisher Scientific) and 1% penicillin/streptomycin (Thermo Fisher Scientific) and grown until 100% confluency. FBs were then stimulated with 10ng/ml TGFβ1 (HEK-293-derived, Peprotech, Rocky Hill, NJ, USA), and with or without DPP4-inhibitor Sitagliptin (10μM) (Thermo Fisher Scientific) or urokinase-inhibitor BC-11 hydrobromide (10μM) (Tocris by Bio-Techne, Bristol, UK) for 24h. Supernatants were removed and medium and inhibitors were resupplied for another 24h. Supernatants were collected and stored at −80°C and cells were lysed in 1x Laemmli Buffer (Bio-Rad Laboratories, Inc., Hercules, CA, USA) for further analysis.

To analyse signaling pathways, FBs were stimulated with TGFβ1 and inhibitors for 1h, and then harvested in 1x Laemmli Buffer with protease inhibitor (cOmplete, MiniProtease Inhibitor Cocktail Tablets, Roche, Basel, Switzerland) and phosphatase inhibitor (Pierce™Phosphatase Inhibitor Mini Tablets, Thermo Scientific, Waltham, MA, USA).

### Western blotting

Primary FBs were lysed in 1x Laemmli Buffer (Bio-Rad Laboratories, Inc.) and loaded on 4-15% SDS-PAGE gels (Bio-Rad Laboratories, Inc.). Proteins were transferred on a nitrocellulose membrane (Bio-Rad Laboratories, Inc.), membranes were blocked in non-fat milk with 0,1% Tween 20 for 1h (Sigma-Aldrich), and incubated with antibodies as indicated in Table S2 at 4°C overnight. After washing, membranes were incubated with horseradish-peroxidase conjugated secondary antibodies as indicated in Table S2 for 1h at room temperature. Signals were developed with SuperSignal West Dura substrate (Thermo Fisher Scientific) and imaged with a Gel Doc XR+ device (Bio-Rad Laboratories, Inc.).

### Proteome profiling of signaling pathways

To analyse signaling pathways, we used a proteome profiler for human phospho-kinases (ARY003C, R&D Systems, Biotechne, Minneapolis, MN, USA) according to the manufacturer’s instructions.

### Enzyme-linked Immunosorbent assay (ELISA)

Human pro-collagen Ia1 ELISA (R&D Systems) and human fibronectin ELISA (R&D Systems) were performed with supernatants of TGFβ 1-stimulated FBs according to the manufacturer’s manual. Absorbance was detected by FluoStar Optima microplate reader (BMG Labtech, Ortenberg, Germany).

## Results

### The single-cell landscape of hypertrophic scars

To elucidate the complex biological processes of scar formation, we performed droplet-based singlecell transcriptome analysis of human hypertrophic scar tissue and healthy skin (Figure 1A). In both samples, Unsupervised Uniform Manifold Approximation and Projection (*UMAP*)-clustering revealed 21 cell clusters, which were further classified as specific cell types by well-established marker genes (Figure S1A), expression patterns of all clusters (Figure S1B), and transcriptional cluster proximity *via* a phylogenetic cluster tree (Figure 1B). We found seven FB clusters, smooth muscle cells and pericytes (SMC/Peri), three clusters of endothelial cells (EC), and lymphatic endothelial cells (LECs), two clusters of T-cells and of dendritic cells (DC), macrophages (Mac), three keratinocyte (KC) clusters, and melanocytes. All cells of specific subsets were clustered together, and skin and scar samples displayed comparable cellular cluster composition (Figure 1C, D). Only cluster FB1 was mainly present in scar tissue. The clusters of skin and scars showed different relative cell number ratios (Figure 1E, F). Whereas FBs represented 40% of all cells in healthy skin, a significant increase (53%) was observed in scar tissue. Similarly, we detected more ECs (16,31%) in scar tissue as compared to normal skin (8,1%). Contrary, the relative numbers of KCs (5,97%) and macrophages (4,42%) in mature hypertrophic scars were significantly reduced compared to skin (18,97% and 7,11%).

**Figure 1:**
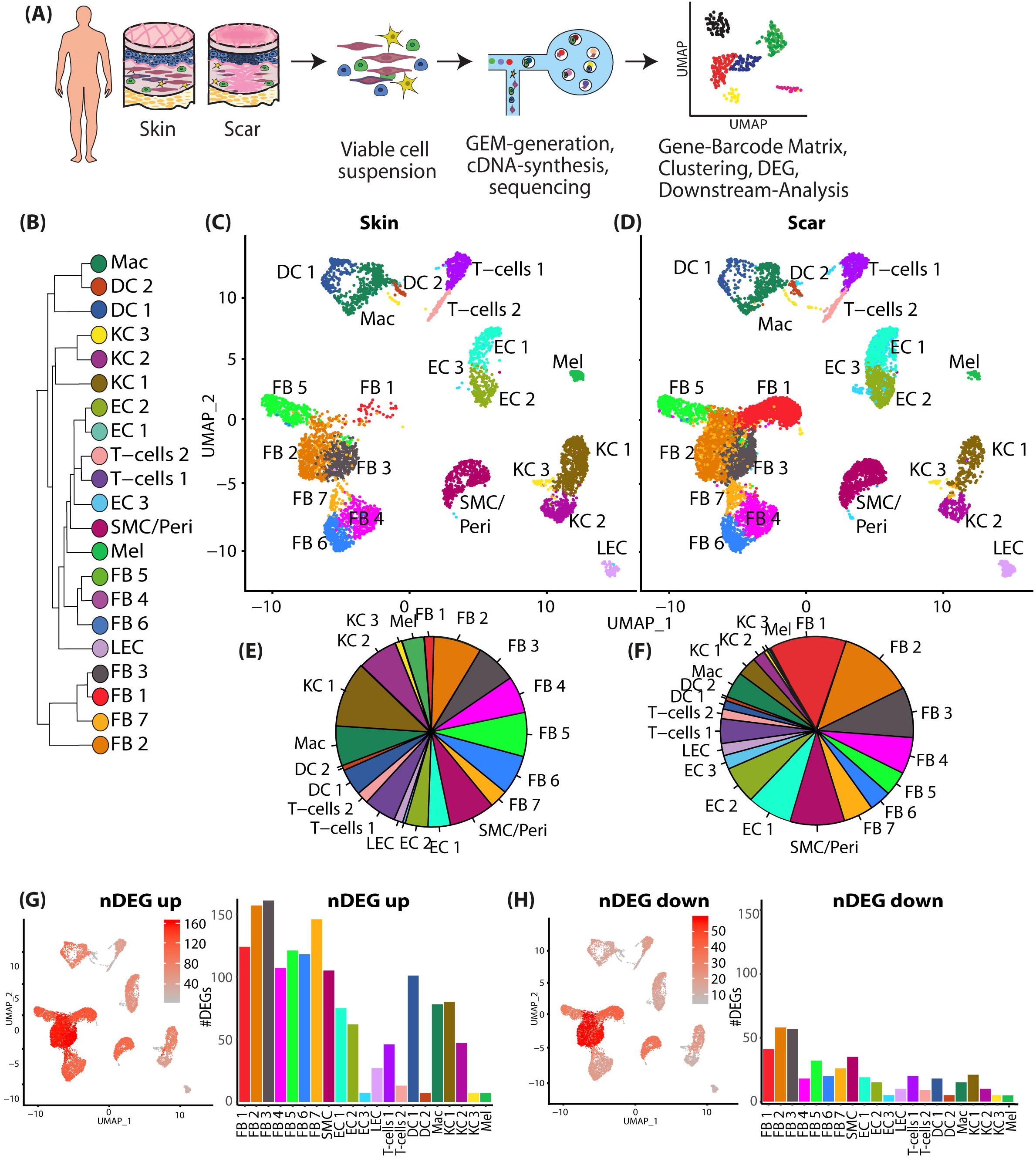
Characterization of human skin and scar samples with scRNAseq identifies specific cell clusters and a distinct fibrotic gene signature. A) Illustration of workflow of scRNAseq in human skin and scar samples. B) Phylogenetic clustertree calculated unsupervised based on UMAP-clustering. C, D) UMAP-plots of human skin and scar samples, split by tissue, after integration of all samples, identifying seven fibroblast clusters (FB1-7), smooth muscle cells and pericytes (SMC/Peri), endothelial cells (EC1+2), lymphatic endothelial cells (LEC), T-cells, macrophages (Mac), dendritic cells (DC1+2), three keratinocyte clusters (KC1-3), and melanocytes (Mel). E, F) Pie charts showing ratios of cell clusters in skin and scars. Feature plots and bar graphs of number of differentially expressed genes (nDEG) per cluster of G) up-and H) downregulated genes. DEGs were calculated per cluster comparing scar versus skin using Wilcoxon rank sum test, including genes with average logarithmic fold change (avglogFC) of > 0.1 or < −0.1 and Bonferroni-adjusted p-value <0.05. Feature plots show projection of nDEG onto the UMAP-plot, color intensity represents nDEG. Bar graphs show absolute numbers of nDEG per cluster, y-axis represents nDEG. UMAP, uniform manifold approximation and projection.

To identify the strongest differences in gene expression between normal skin and scars, we calculated all up- and downregulated genes for FBs, PCs, ECs, T-cells, DCs and KCs (Figure S2). In general, we identified considerably more up- (Figure 1G) than downregulated genes (Figure 1H), and the most abundant differential gene expression of scar compared to skin (number of differentially expressed genes, nDEG) was found in FBs, SMC/pericytes, macrophages, DC1 and KC1 (Figure 1G, H). Genes related to ECM production were mainly overrepresented in FBs, but notably also in PCs and ECs (Figure S2A-C). Several significantly regulated genes with so far undescribed roles in fibrosis and scar formation were found in all cell types (Figure S2A-F). These distinctly regulated genes might provide valuable new candidates to understand and modulate skin scarring.

### The fibrotic gene expression pattern of fibroblasts in hypertrophic scars

Since FBs showed strongest gene regulation in our scRNAseq dataset, and have been considered as the major drivers of skin scarring and an important source for myofibroblasts (Hinz, 2016), we focused our further analysis on differences between FBs of healthy skin and hypertrophic scars (Figure 2).

**Figure 2:**
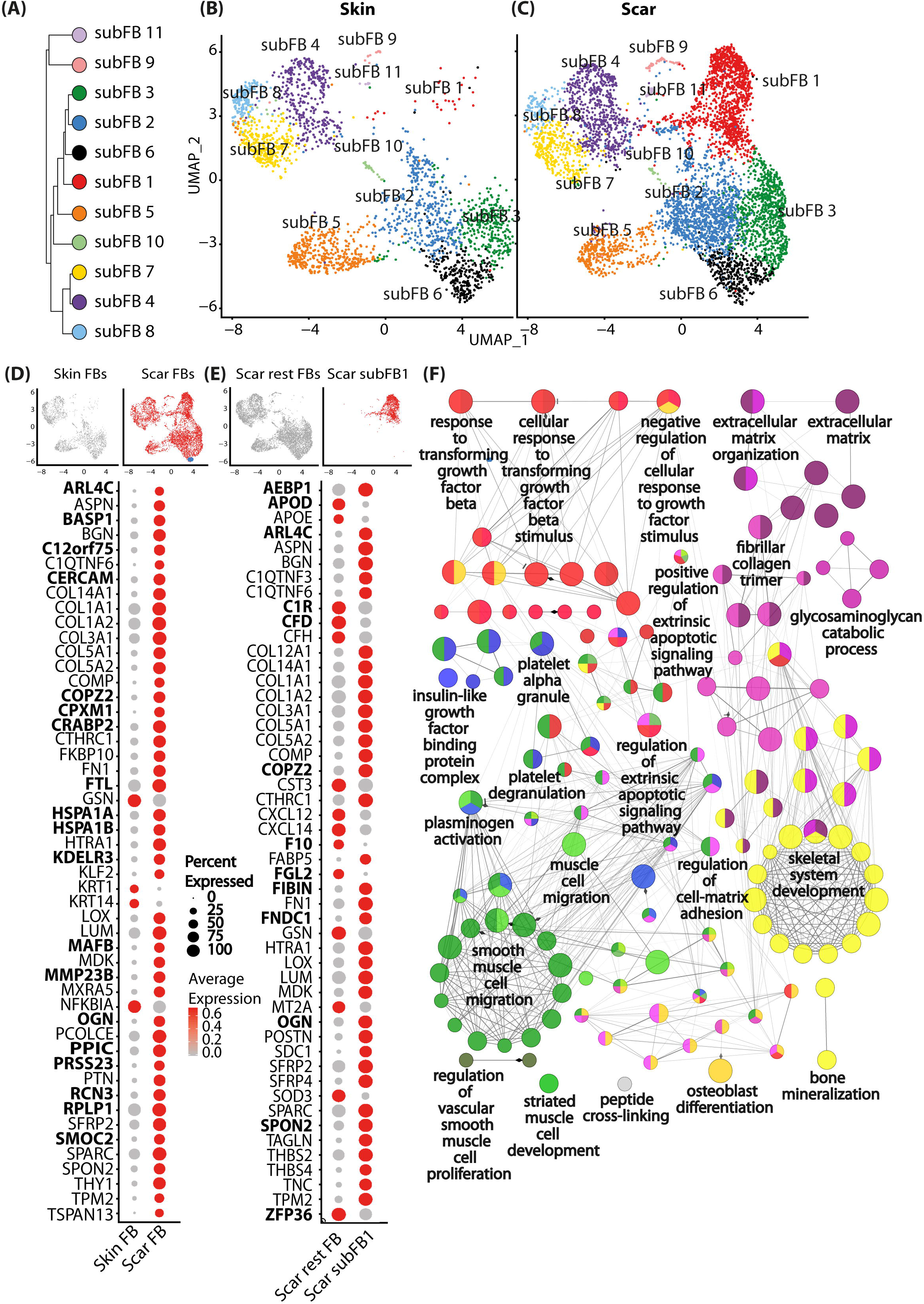
Subset analysis of fibroblasts. A) Phylogenetic clustertree calculated unsupervised based on UMAP-clustering of subsetted fibroblasts only. B, C) UMAP-plots of re-clustered skin and scar fibroblasts, split by tissue, reclustering identified 11 fibroblast clusters (subFB1-11). D) Feature plots illustrating computational basis for dotplots, dotplots of top 50 regulated genes (according to lowest adjusted p-value) comparing scar FBs versus skin FBs. E) Dotplot of top 50 regulated genes (according to lowest adjusted p-value) cluster subFB1 compared to all other scar FBs. F) Gene ontology-term network was calculated based on significantly upregulated (adj. p-val <0.05, avg.logFC >0.1) genes comparing subFB1 to all other scar FBs. Gene list was imputed in ClueGO plug-in in Cytoscape with medium GO-specificity, with GO-term fusion, only significant (*P* value <.05) GO terms are shown. Circle size correlates with *P* value, lines (“edges”) represent functional connection of respective GO terms. Red circles represent association of GO-term with TGFβ-signaling, purple, with extracellular matrix, green, with smooth muscle differentiation, blue, with signaling factors, and yellow with bone formation and -development. UMAP, uniform manifold approximation and projection.

After subsetting and reclustering of all FBs, we identified 11 separate clusters (Figure 2A-C) showing 110 significantly up- and 85 downregulated genes in FBs derived from scar tissue compared to healthy skin. The top 50 differentially up- and downregulated genes are shown in Figure 2D. Interestingly, one FB cluster (FB1) was almost exclusively present in hypertrophic scars, suggesting a specific role in tissue fibrosis. Comparison of FB cluster 1 to all other scar FBs revealed 141 significantly up- and 179 downregulated genes. The top 50 differentially up- and downregulated genes are shown in Figure 2E. Most of the upregulated genes in scar-derived FBs are well-studied in the context of skin scarring and are functionally related to collagens and ECM-modifying genes (Figure S3A, B). However, the interaction on protein level of several genes found here (Figure S3A, B, bold) was hitherto not studied with regard to scar formation.

Analysis of the biological processes associated with differentially regulated genes between FB1 and other FB clusters by gene ontology network analysis revealed a strong association of FB1 with TGFβ-signaling (red circles) and ECM-formation (purple circles) (Figure 2F), further corroborating its role in skin fibrosis. In addition, our analysis indicates a role of FB1 in processes important for several other cell types, including platelets, smooth muscle cells and cells of the skeletal system, suggesting paracrine actions of FB1.

Pseudotime calculation and trajectory construction effectively identifies possible cell fates and time-regulated genes, even when analyzing cells of only one timepoint (Qiu et al, 2017b; Trapnell et al, 2014). Thus, we next sorted human skin and scar FBs along a pseudotime axis and constructed trajectories (Figure 3A, B). The trajectories revealed a division at a certain time point where FBs divided into two branches (Figure 3C). Whereas the majority of FBs preferentially aligned with branch 1 in normal skin (Figure 3D), we observed a significantly longer branch 2 with FBs of hypertrophic scar tissue (Figure 3E). Genes most regulated in a pseudotime-dependent manner in normal skin and hypertrophic scars are shown in Figure 3F and 3G. Interestingly, the collagens *COL1A1, COL1A2* and *COL3A1,* known to contribute to all fibrotic processes, are most upregulated at the end of skin pseudotime, but are not among the most pseudotime-regulated genes in scar (Figure 3F,G). In contrast, COL5A1/2, COL8A1, COL11A1 and COL12A1, dominated the late pseudotimedependent gene expression in scars. The role of these collagens in (hypertrophic) scar is scarcely investigated, and merits further exploration. Together, our trajectory analysis models the temporal dynamics of gene expression in scars and might provide a basis to target respective genes at different stages of scar development.

**Figure 3:**
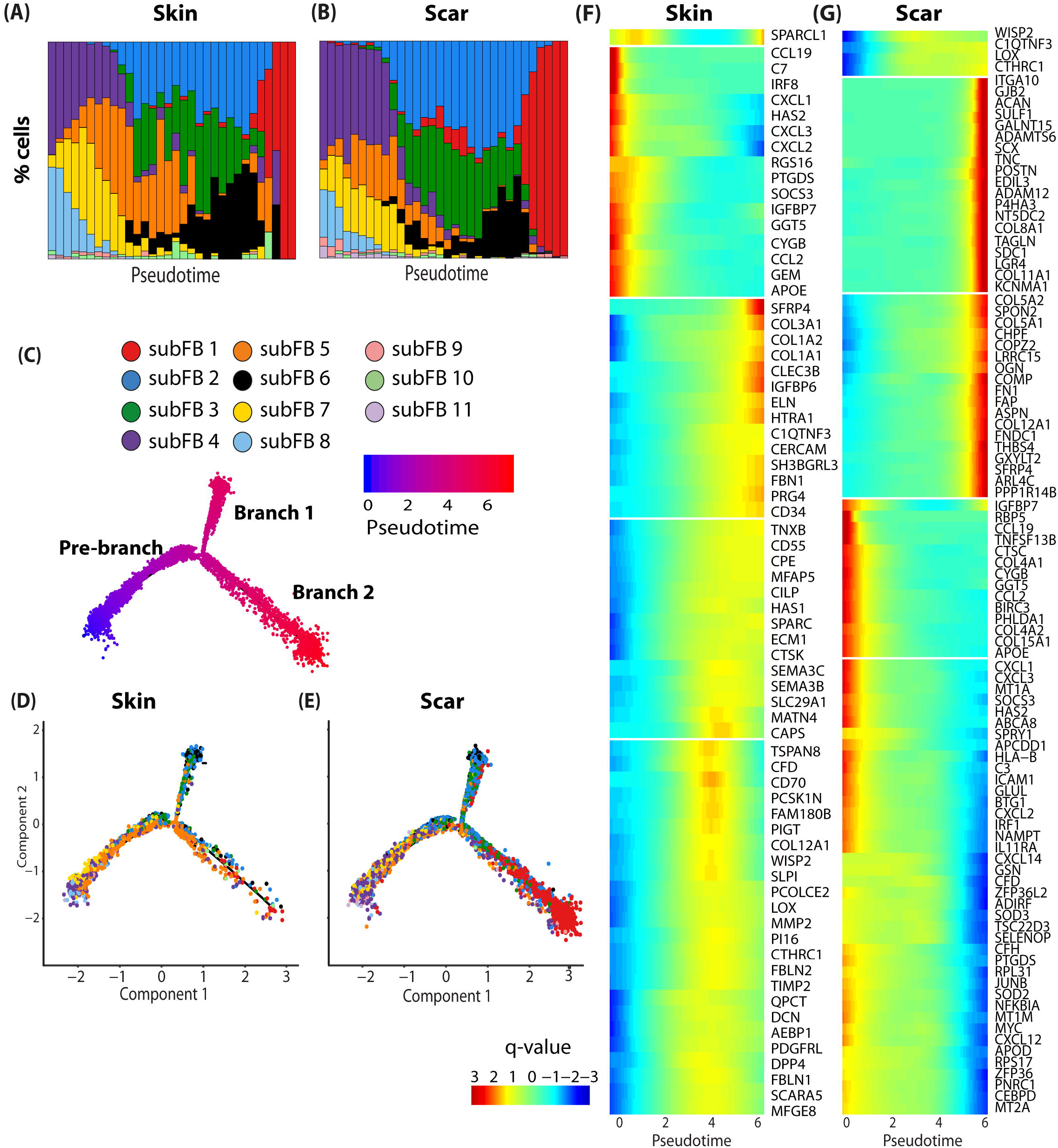
Pseudotime analysis of human scar FBs identifies cell fates and time-regulated genes. A, B) Ordering skin and scar FBs along a pseudotime axis. X-axis, pseudotime. y axis, % of cells in respective pseudotime-bin. Cell trajectory with pre-branch and branches is shown. Color code represents pseudotime progression. C, D) Cell trajectories were calculated based on pseudotime values, split by tissue. E, F) Heatmaps of pseudotime-dependent gene expression in skin and scar. Colors represent q-value, the expression of the respective gene in relation to pseudotime.

### scRNAseq of murine scars identifies genes involved in scar maturation

As our approach so far only gave information on the current state of mature scars, we further investigated mechanisms leading to scar formation and maturation, using a murine full thickness skin wound model. Whereas scar formation and maturation in humans is a long lasting process (Kant et al, 2019), it only takes up to 80 days in rodents (Ferguson & O’Kane, 2004). Hence, we compared scar formation in mice 6 and 8 weeks after wounding (Figure 4A). Analogously to the human dataset, all cells were clustered, and cell types were identified using established marker genes (Figure S4A), expression patterns of all clusters (Figure S4B) and transcriptome proximity of clusters *via* a phylogenetic cluster tree (Figure 4B). All clusters aligned homogeneously, and all major skin cell types were represented at both time points (Figure 4C, D).

**Figure 4:**
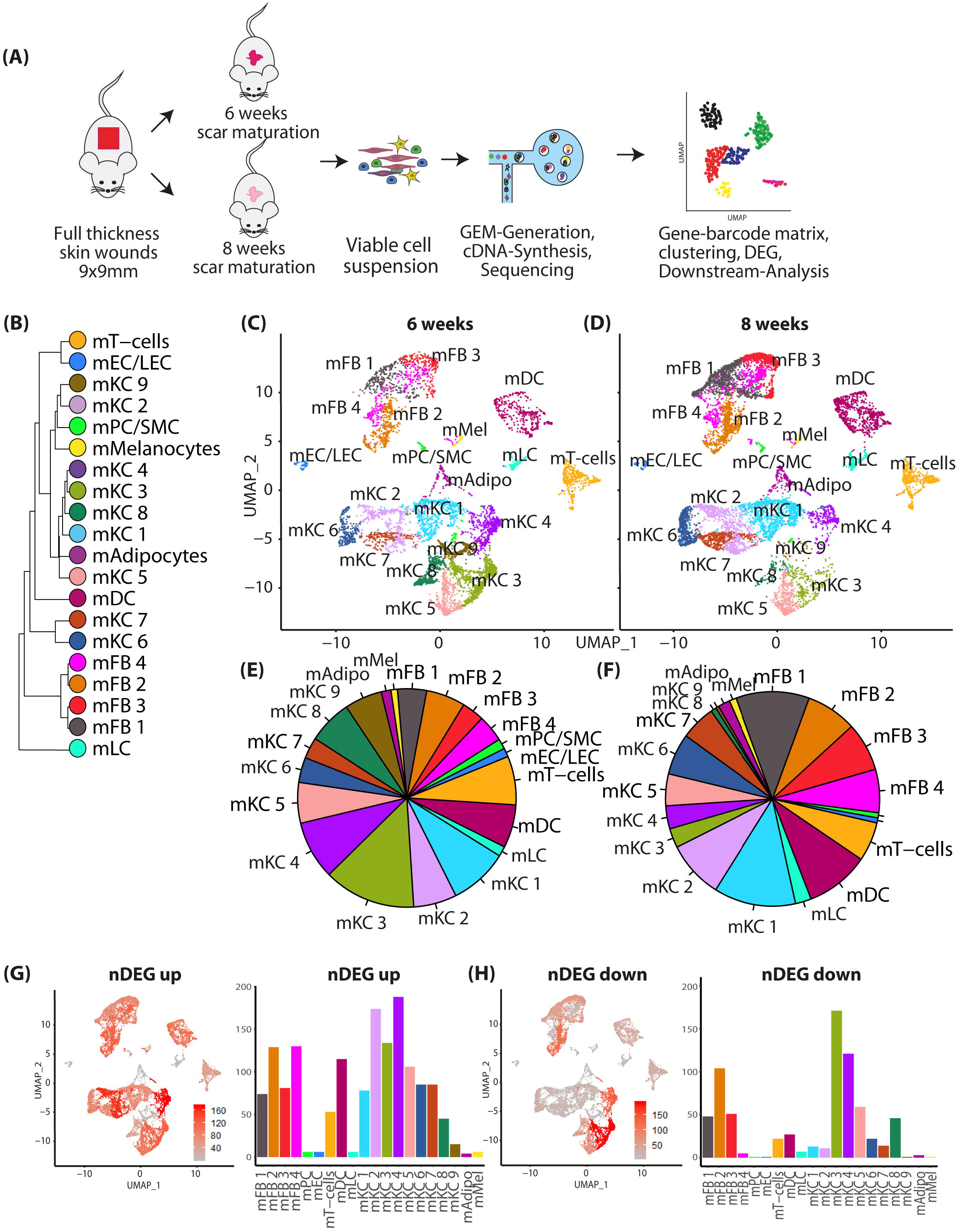
Two-timepoint mouse scar model identifies genes regulated in scar maturation. A) Illustration of workflow of mouse scar model and two-timepoint scRNAseq. Two mice were analyzed per timepoint. B) Phylogenetic clustertree calculated unsupervised based on unsupervised UMAP-clustering. C, D) UMAP-plots of mouse scar tissue, split by timepoint, after integration of all samples, identifying four fibroblast clusters (mFB1-4), smooth muscle cells and pericytes (mPC/SMC), endothelial cells and lymphatic endothelial cells (mEC/LEC), T-cells, dendritic cells (mDC), Langerhans cells (mLC), nine keratinocyte clusters (KC1-9), adipocytes (mAdipo) and melanocytes (Mel). E, F) Pie charts show relative numbers of cells in clusters, split by timepoint. Feature plots and bar graphs of number of differentially expressed genes (nDEG) per cluster of G) up- and H) downregulated genes per cluster. DEGs were calculated per cluster comparing 8 weeks vs 6 weeks old scars using Wilcoxon rank sum test, including genes with average logarithmic fold change (avg_logFC) of > 0.1 or < −0.1 and Bonferroni-adjusted p-value <0.05. Feature plots show projection of nDEG onto the UMAP-plot, color intensity represents nDEG. Bar graphs show absolute number of nDEG per cluster, y-axis represents nDEG. UMAP, uniform manifold approximation and projection.

In accordance with human scar tissue, 8 weeks old mouse scars contained a higher proportion of murine FBs (mFBs) (32,6%) compared to 6 weeks old scars (17,39%), and more DCs (9,6 versus 6,3%). In contrast, less pericytes and endothelial cells (2,8 versus 1,5%) and less keratinocytes (63,3 vs 45%) were present (Figure 4E, F,). We next calculated up- and downregulated genes for FBs, PCs, ECs, T-cells, DCs and KCs, comparing 8 weeks to 6-week-old scars (Figure S5A-F). In contrast to human scars, the highest number of differentially expressed genes was found in mFBs and mKCs (Figure 4G,H), which was most likely due to ongoing epidermal tissue regeneration. Surprisingly, the expression of collagens did not change significantly between 6 and 8 weeks in mFBs, and only mECs showed a slightly increased expression of collagens (Figure S5A-C). However, expression of several matricellular proteins and other ECM-components, e.g. *Fbln1* (Fibulin1), *Ogn* (osteoglycin), *Lum* (Lumican), and *Pcolce* (Procollagen C-Endopeptidase Enhancer) increased in mFBs during scar maturation (Figure S5A). Together, our scRNAseq identified a gene profile specific for scar maturation in mice.

### Serine proteases are strongly upregulated during scar maturation

To identify genes that are crucial for scar maturation, we next compared our human scar data set with genes upregulated in mouse scars 8 weeks after wounding in comparison to mouse scars 6 weeks after wounding (Figure 5A). While in both data sets only one gene *(LEPR)* was downregulated, 16 genes were mutually upregulated (Figure 5B-D). Stunningly, 5 of these genes *(AEBP1, DPP4, HTRA1, PLAU* and *PRSS23)* were members of the superfamily of serine proteases (Figure 5B, C). All five serine proteases were upregulated in scRNAseq in human scar tissue, particularly in FBs, but also in other cell types (Figure 5E-I). *AEBP1* and *PRSS23* expression also increased in ECs and melanocytes, *HTRA1* in ECs and KC3, and *PLAU* in DCs. Several additional serine proteases, *HTRA3* (High-Temperature Requirement A Serine Peptidase 3), *DPP7* (Dipeptidyl-Peptidase 7), *FAP* (Fibroblast activation protein alpha), were upregulated in human scars (Figure S6), and also showed a trend in mouse scars. Analysis of theses serine proteases by pseudotime trajectories in human FBs revealed that their expression mainly increased over time and *AEBP1* and *HTRA1* significantly enriched at the end of branch 2 (Figure 6A). Together, these data suggest a major role of serine proteases in scar formation and/or maturation.

**Figure 5:**
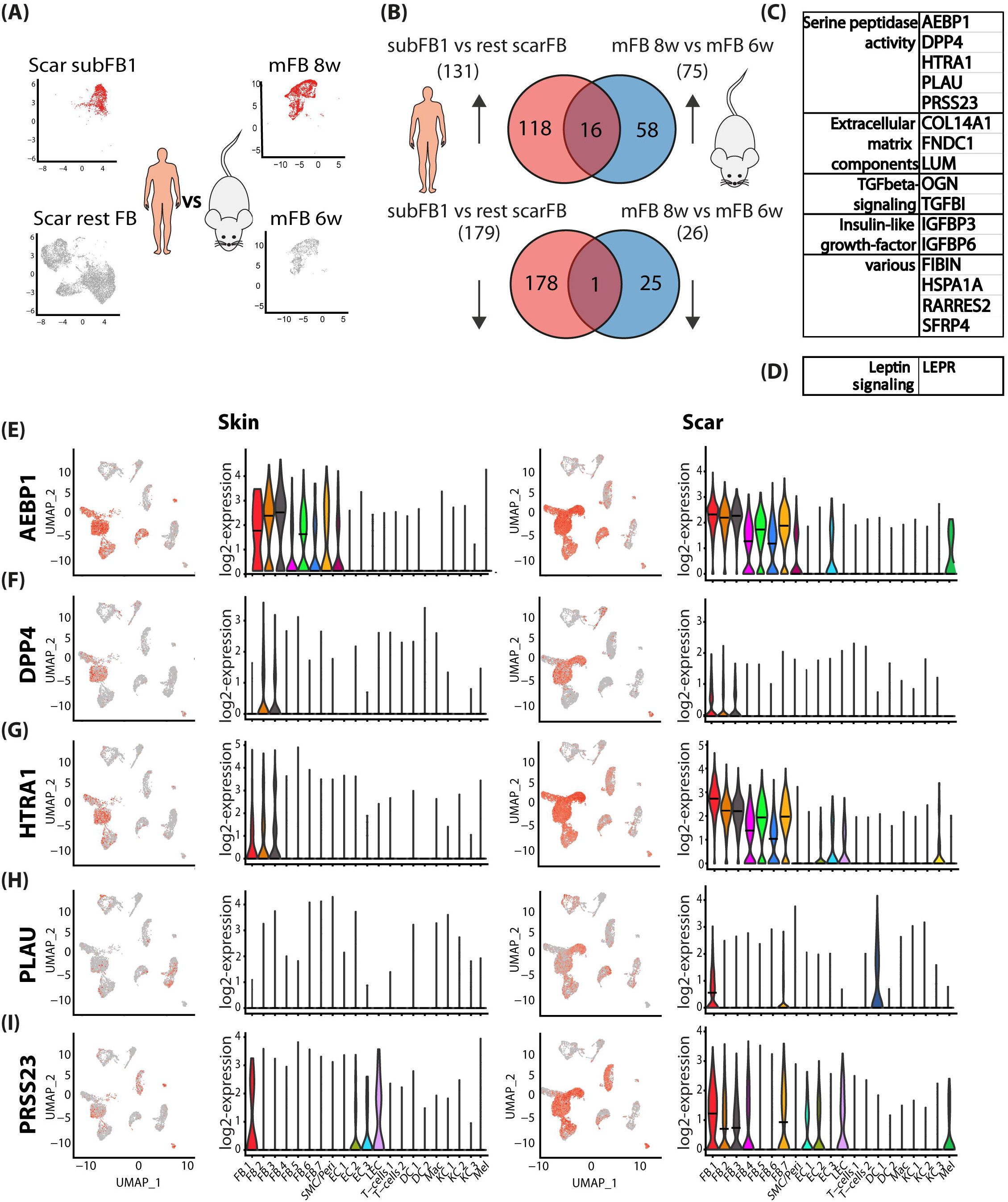
Comparing human scar gene expression and mouse scar maturation identifies mutual drivers of skin fibrosis. A) Illustration of computational basis for comparison human and mouse. Human cluster subFB1 vs rest scar FBs significantly (adj. p-value<0,05) regulated genes were compared with mouse scar FBs 8 weeks vs 6 weeks significantly regulated genes. B) Venn diagrams of human and mouse up- (upper panel) and down- (lower panel) regulated genes. C) Table of mouse and human mutually up and D) downregulated genes. E-I) Feature plots and violin plots of serine proteases in human skin and scar. *AEBP1* (adipocyte enhancer binding protein 1), *DPP4* (dipeptidyl-peptidase 4), *HTRA1* (High-Temperature Requirement A Serine Peptidase 1), *PLAU*(urokinase), *PRSS23* (Serine protease 23). In violin plots, dots represent individual cells, *y*-axis represents log2 fold change of the normalized genes and log-transformed single-cell expression. Vertical lines in violin plots represent maximum expression, shape of each violin represents all results, and width of each violin represents frequency of cells at the respective expression level. In feature plots, normalized log expression of the respective gene is mapped onto the UMAP-plot. Color intensity indicates level of gene expressions. UMAP, uniform manifold approximation and projection.

**Figure 6:**
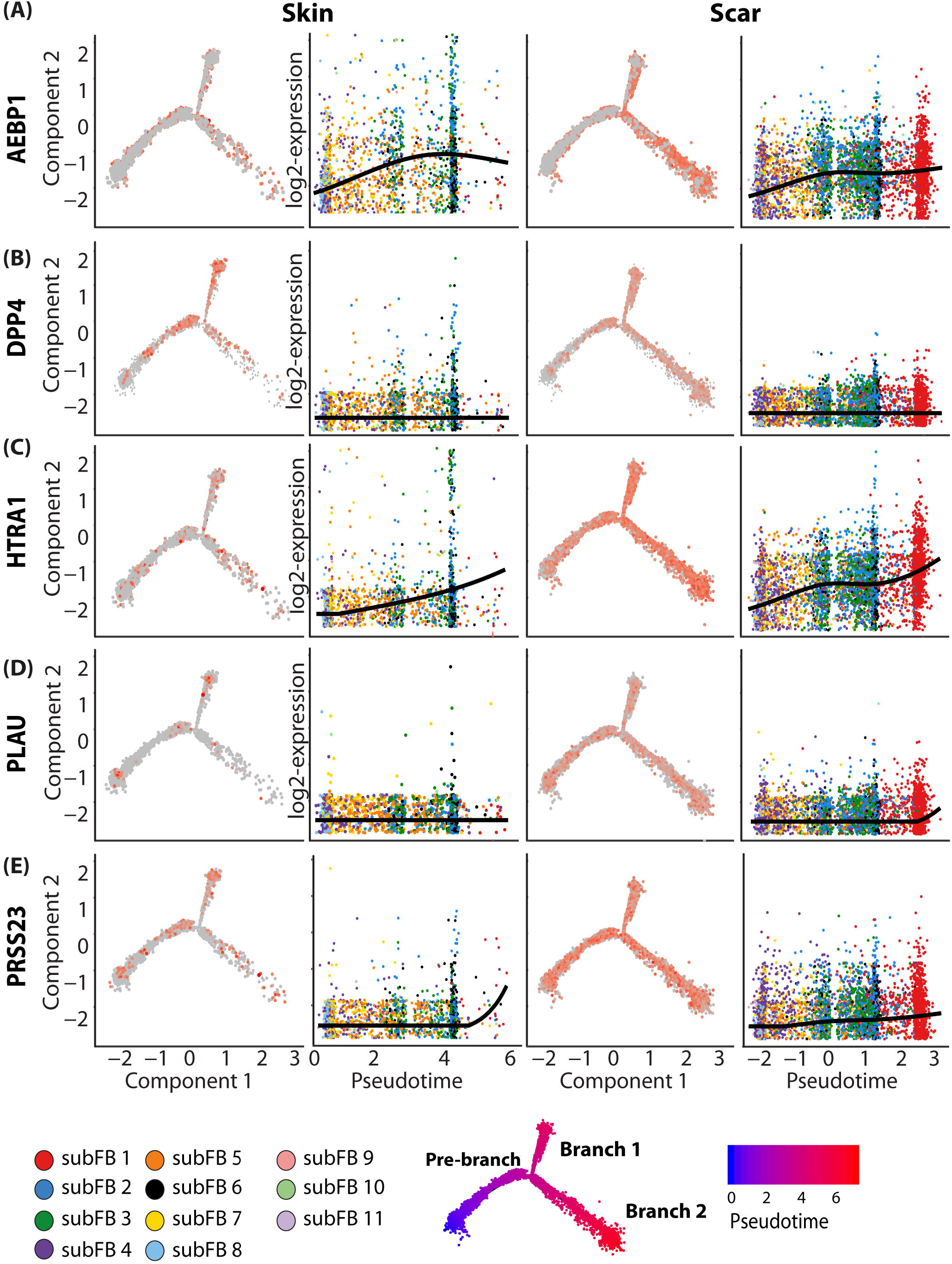
Pseudotime analysis corroborates the putative role of serine proteases as drivers of scar maturation. A-E) Trajectory plots and pseudotime plots of Serine proteases in skin and scar FBs. *AEBP1* (adipocyte enhancer binding protein 1), *DPP4* (dipeptidyl-peptidase 4), *HTRA1* (High-Temperature Requirement A Serine Peptidase 1), *PLAU* (urokinase), *PRSS23* (Serine protease 23). In trajectory plots, normalized log expressions are plotted on the trajectories, split by skin and scar. In pseudotime plots, normalized log expressions are plotted against the pseudotime axis, and a spline curve represents expression dynamics over pseudotime. Y-axis, normalized log expression of respective gene, x-axis, pseudotime.

### The serine proteases DPP4 and urokinase regulate TGFβ1-mediated myofibroblasts differentiation and ECM over-production

We next wanted to investigate the contribution of the identified serine proteases to scar formation. Since specific inhibitors are commercially available only for DPP4 and urokinase, we focused our further functional studies on these two serine proteases. First, we corroborated our scRNAsec data by analyzing RNA and protein expression of DPP4 and urokinase (*PLAU*) using in-situ hybridization (Figure 7A, B), and immunofluorescence stainings (Figure 7C, D), respectively. Both methods showed significantly higher expression of DPP4 and PLAU in hypertrophic scars as compared to healthy skin. In addition to dermal FBs, *PLAU* was also induced in the epidermis of scar tissue. As we have previously identified a strong upregulation of *PLAU* in scars in DC 1 (Figure 5H), the positive cells in the epidermis most likely represent Langerhans cells.

**Figure 7:**
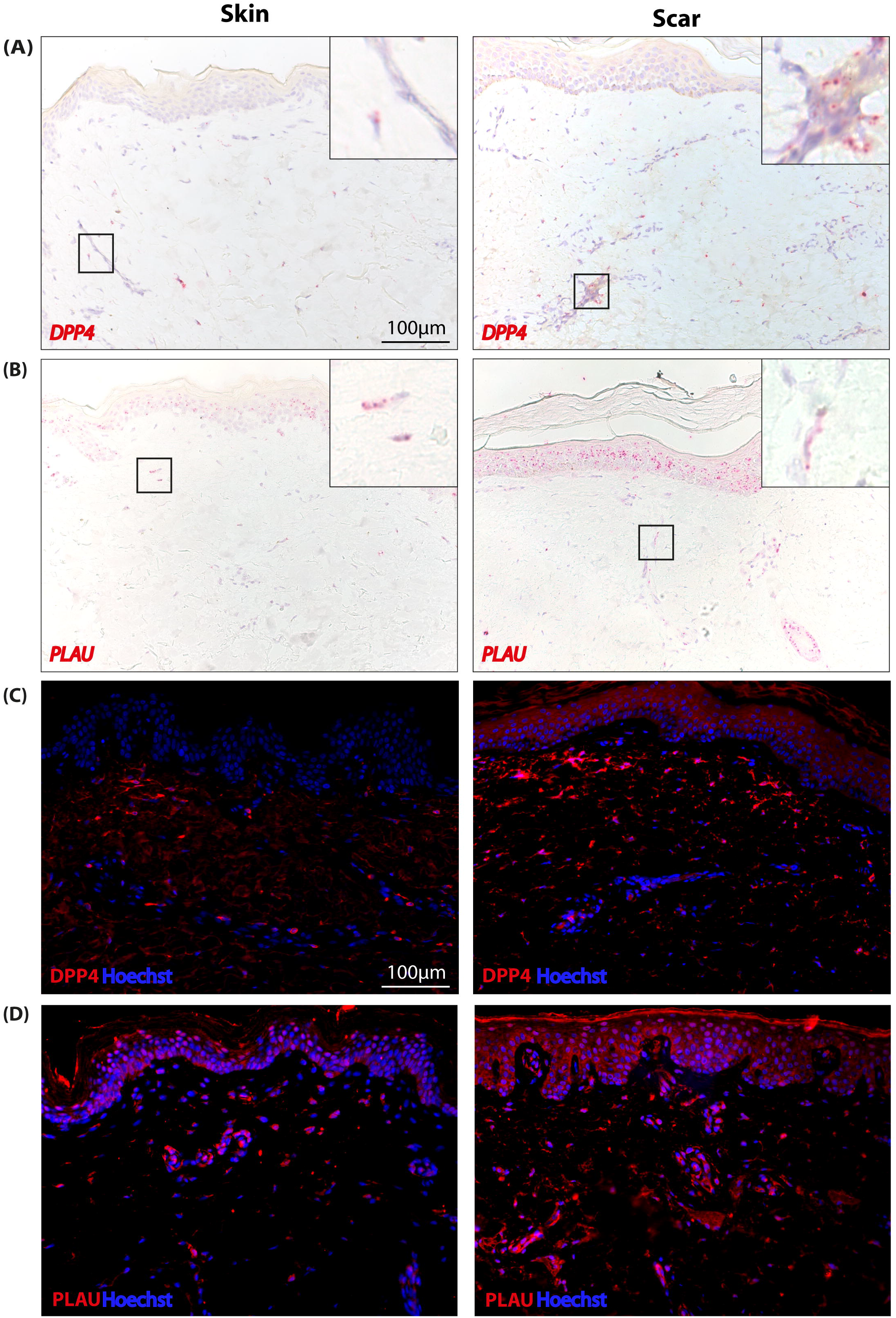
*In-situ* hybridization and immunofluorescence staining confirm elevated expression of *PLAU* and *DPP4* in human skin and scar. A, B) RNAScope in-hybridization of skin and scar tissue with *DPP4-* and *PLAU-probes.* Red dots indicate single mRNA molecules. Inserts show high magnification micrographs. C, D) Immunofluorescent staining of DPP4 and PLAU in human skin and scar tissues.

As TGFβ1 is one of the key inducers of scarring and tissue fibrosis, causing differentiation of FBs to pro-fibrotic myofibroblasts (Carthy, 2018; Lodyga & Hinz, 2019; Roberts et al, 1986; Schuliga et al, 2017), we hypothesized that the serine proteases interact with TGFβ-signaling. To test this, we stimulated primary FBs from healthy abdominal human skin with active TGFβ1, which induces expression of alpha-smooth muscle actin (αSMA), a marker for myofibroblasts. Addition of the DPP4 inhibitor Sitagliptin and the urokinase inhibitor BC11 almost completely abolished the TGFβ1-mediated myofibroblast-differentiation as shown by αSMA Western blot (Figure 8A, B). Moreover, Sitagliptin and to an even greater extent BC-11 attenuated TGFβ1-induced overproduction of the ECM-proteins Col1a1 (Figure 8C, D), and fibronectin (Figure 8E, F) by FBs. These results suggest that serine proteases are involved in the TGFβ1-induced myofibroblast differentiation.

**Figure 8:**
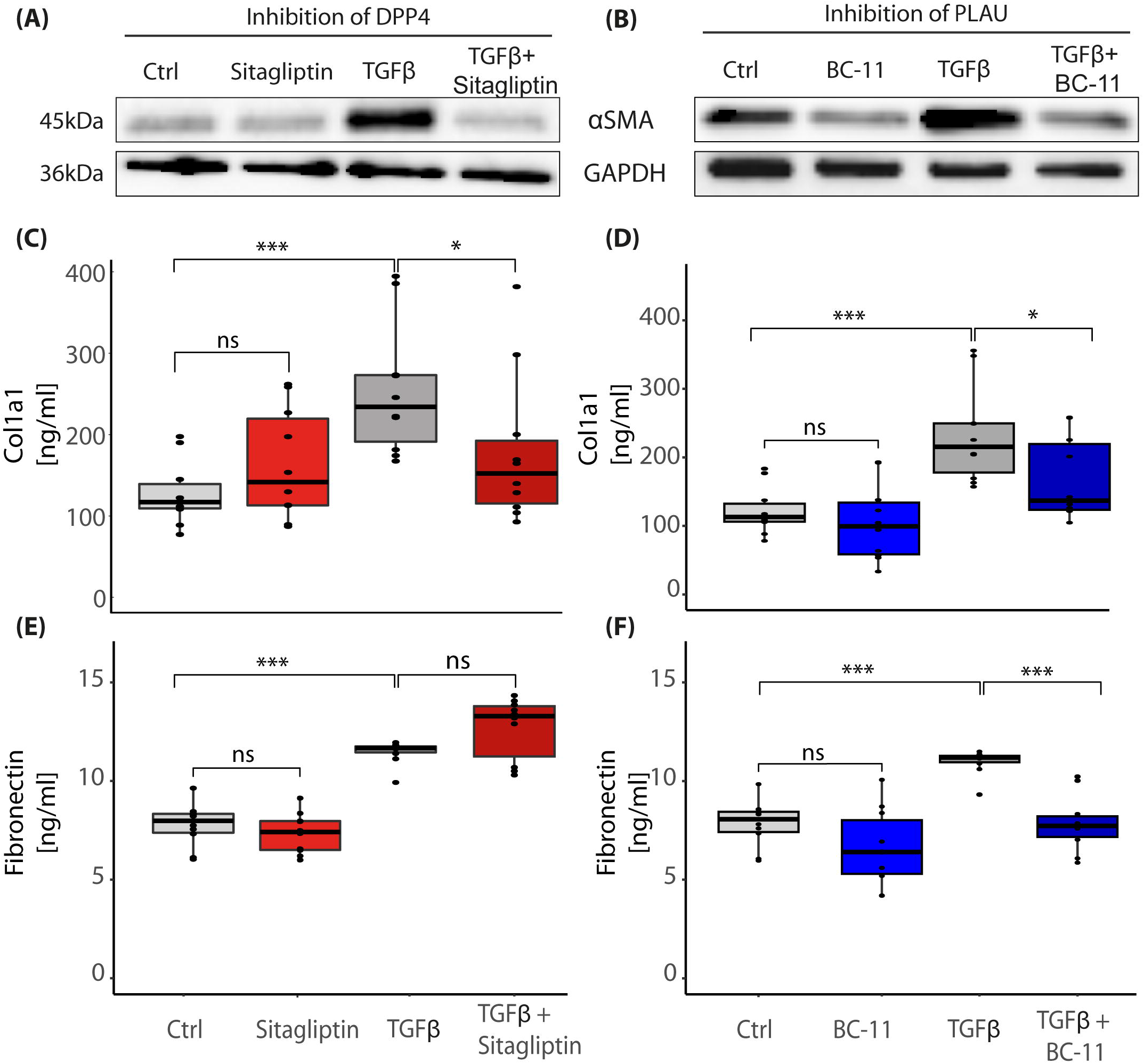
Pharmacological inhibition of DPP4 or urokinase prevents TGFβ-induced myofibroblast-differentiation. A,B) Western blot of primary FBs stimulated with active TGFβ1 for 24h to differentiate FBs into alpha-smooth muscle actin-expressing (αSMA) myofibroblasts. Myofibroblast differentiation inhibited with A) DPP4-inhibitor Sitagliptin or B) urokinase-inhibitor BC-11. C, D) Collagen I or E, F) fibronectin in supernatants of stimulated primary skin FBs, detected by Enzyme-linked Immunosorbent Assay (ELISA). Whiskers represent range maximum and minimum values with < 1.5 interquartile range, boxes represent 25^th^ −75^th^ quartiles, line represents mean. Statistical significance was tested using one-way ANOVA with Tukey post-test. NS p>0.05, *p<0.05, **p<0.01, ***p<0.001. Experiments were performed in duplicates of five donors each.

### The serine proteases DPP4 and urokinase do not interfere with canonical TGFβ1-signaling pathways

To investigate whether the serine protease inhibitors interfere with TGFβ1 signaling, we investigated the TGFβ1-induced signaling pathways SMAD2 and ERK1/2 (Hata & Chen, 2016). While SMAD2 and ERK1/2 phosphorylation was readily induced upon stimulation with TGFβ1, addition of the inhibitors showed no effects (Figure 9A). In contrast, SMAD1, 5 and 9 showed no activation after TGFβ1 stimulation (Figure 9A). To further identify other signaling pathways that might be involved in the action of the serine protease inhibitors, we used a signaling proteome profiler, showing that none of the signaling molecules were inhibited by the inhibitors (Figure 9B). Interestingly, the GSKα/β-pathway, known to attenuate fibrotic processes in the heart (Lal et al, 2014) was significantly activated by BC-11 (Figure 9B, Figure S7), suggesting a counter-regulatory action. Together, these data suggest that sitagliptin and BC-11 do not interfere with canonical or known non-canonical TGFβ1 signaling.

**Figure 9:**
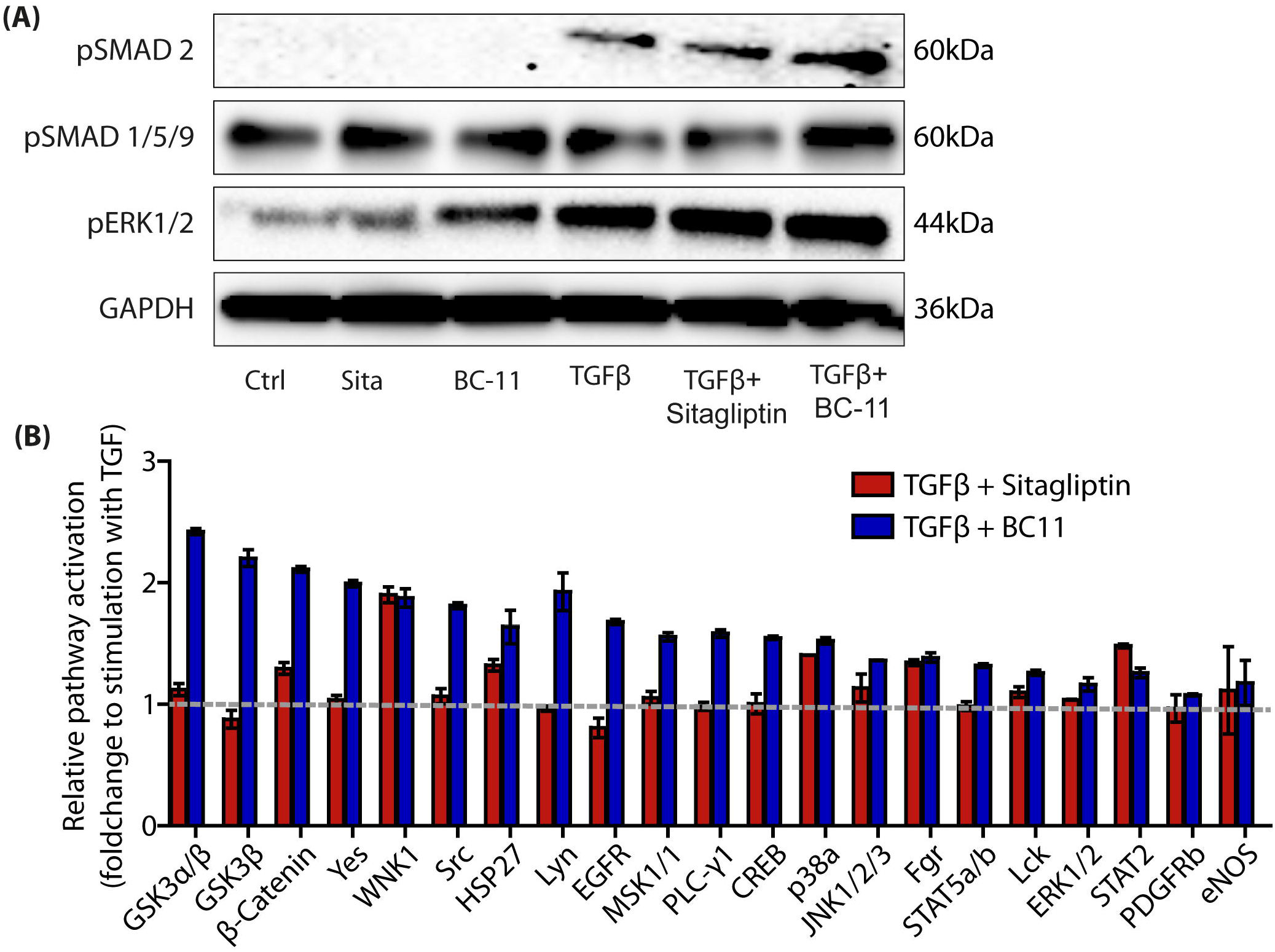
Signaling pathway activation after TGFβ-stimulation and DPP4/PLAU inhibition. E) Western blot of primary FB lysate stimulated with active TGFβ1 for 1h and analysis of canonical TGFβ1 signaling pathways. B) Analysis of non-canonical TGFβ1 signaling pathways and kinase pathways detected by proteome profiler from lysates of primary FBs after 1h stimulation with TGFβ1 alone, TGFβ1 with sitagliptin, and TGFβ1 with BC-11. Bars represent fold change compared to stimulation with TGFβ1 only, marked by dotted line.

## Discussion

Although skin fibrosis has been extensively studied, key mechanisms leading to the development of hypertrophic scars are still not well understood. In addition, treatment options to prevent or treat (hypertrophic) scars are still scarce (Sidgwick et al, 2015) and not exceptionally effective. In the present study, we used scRNAseq to elucidate the genetic landscape of hypertrophic scar tissue at a hitherto unmet single cell resolution. We found numerous regulated genes not yet investigated in scar formation, and identified serine proteases as regulators of TGFβ-induced myofibroblast differentiation.

As expected, our scRNAseq analysis confirmed a plethora of previous studies, but also identified numerous genes, which have so far not been described in the context of skin scarring or tissue fibrosis. For example, the cytokines *MDK* (midkine) and *PTN* (pleiotrophin), both involved in cell growth, migration, and angiogenesis (Muramatsu, 2010), were strongly upregulated in scar FBs. In contrast, *SOD2/3* (superoxide dismutase 2/3), an enzyme controlling the release of reactive oxygen species (ROS), hence acting as important antioxidant (Miao & St Clair, 2009), was strongly downregulated in scar FBs. Interestingly, failure of ROS-scavenging has already been shown to contribute to hypertrophic scar formation (Carney et al, 2019). Another interesting and significantly down-regulated gene in scars was *SFN* (stratifin). As stratifin has been identified as potent collagenasestimulating factor in FBs, its down-regulation in scars suggests a contribution to the maintenance and/or progression of the fibrotic phenotype by preventing matrix degradation. These and many other novel factors identified in our study could be important, decisive molecules for the development and/or maturation of hypertrophic scars. Thus, our study has built a basis for future studies describing the role of these molecules in skin scarring and tissue fibrosis.Our combined study of human mature hypertrophic scars and scar maturation in mice identified a group of serine proteases as key player for scar development and maturation. Although, DPP4-positive FBs have previously been identified as the main source of ECM production in the skin (Vorstandlechner et al, 2020), and urokinase has been shown to be involved in lung fibrosis (Schuliga et al, 2017), their roles in myofibroblast differentiation and production of ECM are still unclear. Our finding that addition of specific DPP4 and urokinase inhibitors to TGFβ1-stimulated FBs almost completely inhibited myofibroblast differentiation and up-regulation of matrix proteins was striking. Sitagliptin, the here used DPP4 inhibitor, is an effective drug widely used for the treatment of diabetes mellitus (Juillerat-Jeanneret, 2014). Based on our study, it might be very interesting to systemically evaluate differences in scar formation and scar quality of diabetic patients treated with either gliptins or other drugs. Furthermore, topical application of gliptins, during the phase of tissue remodeling after full wound closure, might prevent (hypertrophic) skin scarring. As gliptins are already approved for clinical use, an off-label topical application would be a promising step forward to test its efficacy on skin scarring *in vivo.* Interestingly, the urokinase inhibitor BC-11 showed an even stronger effect on inhibiting TGFβ1-induced myofibroblast differentiation and matrix production. However, so far it has only been used *in vitro,* and further *in vivo* testing for efficacy and safety is still required. Inhibition of urokinase to attenuate tissue fibrosis *per se* might appear counterintuitive, as urokinase facilitates fibrinolysis and regulates ECM-turnover, eliciting anti-fibrotic action (Behrendt, 2004). However, literature on urokinase inhibitors and fibrosis is contradictory. The best investigated endogenous urokinase-inhibitor, plasminogen activator inhibitor-1 (PAI-1, *SERPINE1),* was found to cause excessive matrix deposition after injury (Rabieian et al, 2018). By contrast, and in line with our results, inhibition of urokinase by PAI-1 suppressed profibrotic response in FBs from fibrotic lungs and prevented cardiac fibrosis in mice (Gupta & Donahue, 2017). Therefore, our study suggests the use of urokinase inhibitor BC-11 as a possible new therapeutic strategy for the treatment of skin scars. Further studies are necessary to fully elucidate its efficacy *in vivo*.

Strikingly, our analyses revealed no influence of the inhibitors on the canonical TGFβ1 signaling pathway. As to the anti-fibrotic action of sitagliptin, we have no mechanism to offer yet. Although DPP4 inhibition has previously been demonstrated to directly inhibit canonical TGFβ signaling *via* Smad2 in renal fibrosis (Wang et al, 2018) and TGFβ-mediated myoFB-differentiation by interfering with ERK-signaling (Thielitz et al, 2008), we were not able to confirm these mechanisms in skin FBs. Possible explanations for alternative the anti-fibrotic action of sitagliptin could be interactions with CXCL12/SDF1 (Stromal derived factor 1), a substrate of DPP4 (Elmansi et al, 2019) which was found to promote scar formation (Nishiguchi et al, 2018), or with components of the ECM, such as fibronectin (Elmansi et al, 2019), which have been shown to bind latent TGFβ (Schuliga et al, 2017), or DPP4 induced cleavage of growth factors or receptors (Elmansi et al, 2019). Deciphering these interactions will be the scope of further mechanistic studies. Regarding BC-11, we found a significant activation of GSK3α/β in TGFβ1-stimulated FBs. Since GSK3β was previously found to interact with WNT/β-catenin signaling (Guo et al, 2017; Vallée et al, 2017), and deletion of GSK-3β induced a pro-fibrotic myofibroblast phenotype in isolated cardiac FBs in mice (Lal et al, 2014), the activation of GSK3α/β suggests a counter-regulation of TGFβ1-signaling. It is therefore conceivable that BC-11, at least partially, exerts its anti-fibrotic action *via* activation of GSK3α/β.

Together, our study provides a genetic landscape of hypertrophic scars which is the basis for further investigations on novel genes and fibrotic processes hitherto not studied in the context of skin scarring. In addition, we identified a group of serine proteases involved in scar formation and/or maturation and further *in vivo* studies will demonstrate the efficacy of serine protease inhibitors on the prevention of scar development and on the improvement of already existing scars.

**LiteraturePrimary Sources**

**Secondary Sources**

## Supplementary figure legends

**Figure S1: Identification of cell types by marker genes and marker gene expression patterns in human skin and scar**

A) Feature Plots of cluster markers *KRT1* (Keratin1) for spinous and granular keratinocytes (KCs), *KRT5* (Keratin 5) for basal KCs, *COL1A1* (collagen I alpha 1) for fibroblasts (FBs), *ACTA2* (smooth muscle actin) for smooth muscle cells and myofibroblasts, *RGS5* (Regulator Of G Protein Signaling 5) for pericytes, *SELE* (E-selectin) for endothelial cells, *LYVE1* (Lymphatic Vessel Endothelial Hyaluronan Receptor 1) for lymphatic endothelial cells, *CD3D* (cluster of differentiation 3D) for T-cells, *ITGAX* (Integrin Subunit Alpha X, CD11C) and *CD1A* for dendritic cells, *AIF1* (Allograft Inflammatory Factor 1) for macrophages, and *MLANA (*Melan-A) for melanocytes. In feature plots, normalized log expression of the respective gene is mapped onto the UMAP-plot. B) Heatmap of top 10 clustermarker (differentially upregulated genes of each cluster compared to the rest of the dataset). Heatmap shows scaled expression values for genes, rows represent genes, columns represent individual cells. DEGs were calculated per cluster comparing scar versus skin using Wilcoxon rank sum test, including genes with average logarithmic fold change (avglogFC) of > 0.1 or < −0.1 and Bonferroni-adjusted p-value <0.05. Feature plot shows projection of nDEG onto the UMAP-plot, color intensity represents nDEG. UMAP, uniform manifold approximation and projection.

**Figure S2: Top 50 regulated genes per cell group in scar compared to skin.**

In cellgroups, i.e. A) fibroblasts (FBs), B) smooth muscle cells and pericytes (SMC/PCs), C) endothelial cells (ECs), D) T-cells, E) dendritic cells (DCs), and keratinocytes (KCs), differentially expressed genes (DEGs) were calculated comparing scar versus skin using Wilcoxon rank sum test, including genes with average logarithmic fold change (avglogFC) of > 0.1 or < −0.1 and Bonferroni-adjusted p-value <0.05. For each cell group, top 50 DEGs according to lowest adjusted p-value are displayed, split by skin and scar. Dot size represents percent of cells expressing the respective gene, color correlates with average expression.

**Figure S3: Gene interaction networks of upregulated genes in FBs**

STRING networks of gene/protein interactions from A) top 50 regulated genes (according to lowest adjusted p-value) comparing skin FBs versus scar FBs and B) top 50 regulated genes (according to lowest adjusted p-value) cluster subFB1 to all other scar FBs. Lines indicate protein interaction with evidence level according to legend. Bold gene names indicate protein with no hitherto described relation to skin scarring.

**Figure S4: Identification of cell types by marker genes and gene expression patterns in mouse scars**

A) Feature Plots of cluster markers *Krt1* (Keratin1) for spinous and granular keratinocytes (KCs), *Krt5* (Keratin 5) for basal KCs, *Mki67* (Marker Of Proliferation Ki-67) for proliferating cells, *Col1a1* (collagen I alpha 1) for fibroblasts; *Pecam* (Platelet And Endothelial Cell Adhesion Molecule 1) for endothelial cells, *Lyve1* (Lymphatic Vessel Endothelial Hyaluronan Receptor 1) for lymphatic endothelial cells, *Acta2* (smooth muscle actin) for smooth muscle cells and myofibroblasts, *Rgs5* (Regulator Of G Protein Signaling 5) for pericytes, *Cd3d* (cluster of differentiation 3D) for T-cells, *Cd14* for dendritic cells, *Cd207* (Langerin) for Langerhans cells, and *Pmel* (Premelanosome Protein) for melanocytes. In feature plots, normalized log expression of the respective gene is mapped onto the UMAP-plot.

B) heatmap of top 10 upregulated clustermarker (differentially expressed genes of each cluster compared to the rest of the dataset). Heatmap showing scaled expression values for genes, rows represent genes, columns represent individual cells.

**Figure S5: Top 50 regulated genes per cell group in 6 weeks and 8 weeks old murine scars.**

In A) fibroblasts (FBs), B) smooth muscle cells and pericytes (SMC/PCs), C) endothelial cells (ECs), D) T-cells, E) dendritic cells (DCs), and keratinocytes (KCs), differentially expressed genes (DEGs) were calculated comparing 8 weeks versus 6 weeks old mouse scars using Wilcoxon rank sum test, including genes with average logarithmic fold change (avglogFC) of > 0.1 or < −0.1 and Bonferroni-adjusted p-value <0.05. For each cellgroup, top 50 DEGs according to lowest adjusted p-value are displayed, split by timepoint. Dot size represents percent of cells expressing the respective gene, color correlates with average expression.

**Figure S6: Feature Plots and Violin plots of serine proteases**

A-C) Feature plots and violin plots of serine proteases in human skin and scar. *DPP7* (dipeptidyl-peptidase 7), *FAP* (Fibroblast Activation Protein Alpha), *HTRA3* (High-Temperature Requirement A Serine Peptidase 3). In violin plots, dots represent individual cells, *y*-axis represents log2 fold change of the normalized genes and log-transformed single-cell expression. Vertical lines in violin plots represent maximum expression, shape of each violin represents all results, and width of each violin represents frequency of respective expression level. In feature plots, normalized log expression of the respective gene is mapped onto the UMAP-plot. Color intensity indicates level of gene expressions. UMAP, uniform manifold approximation and projection.

**Figure S7: A proteome profiler-assisted identification of signaling pathways after stimulation with TGFβ and serine protease inhibitors.**

Proteome profiler analysis of signaling pathways of primary human skin FBs stimulated with A) TGFβ1, B) TGFβ1 and DPP4-inhibitor Sitagliptin, and C) TGFβ1 and PLAU-inhibitor BC-11. D) Legend table for proteome profiler data points.

## Acknowledgements

This research project was financed in part by the FFG Grant “APOSEC” (852748 and 862068; 2015-2019), by the Vienna Business Agency “APOSEC to clinic,” (ID 2343727, 2018-2020), and by the Aposcience AG under group leader HJA. MM was funded by the Sparkling Science Program of the Austrian Federal Ministry of Education, Science and Research (SPA06/055). We thank HPH for his belief in this private-public partnership to augment patients’ health. We thank the Biomedical Sequencing Facility, Center for Molecular Medicine, Vienna, Austria, for the processing and sequencing of our samples. The authors acknowledge the core facilities of the Medical University of Vienna, a member of Vienna Life Science Instruments.

## Author contributions

MM, HJA, ET and VV provided study conception and design; WH and CR provided patient sample material; HJA and MM acquired funding; VV, DC, YC and BG conducted experiments and prepared samples; VV performed data analysis, visualization and figure design; VV, ML, ET, and MM participated in data interpretation; VV, ML and MM drafted the manuscript. All authors reviewed the manuscript.

## Uncategorized References

Anthonissen M, Daly D, Janssens T, Van den Kerckhove E (2016) The effects of conservative treatments on burn scars: A systematic review. Burns: journal of the International Society for Burn Injuries 42: 508–518

Aroor AR, Habibi J, Kandikattu HK, Garro-Kacher M, Barron B, Chen D, Hayden MR, Whaley-Connell A, Bender SB, Klein T, Padilla J, Sowers JR, Chandrasekar B, DeMarco VG (2017) Dipeptidyl peptidase-4 (DPP-4) inhibition with linagliptin reduces western diet-induced myocardial TRAF3IP2 expression, inflammation and fibrosis in female mice. Cardiovascular diabetology 16: 61

Bao Y, Xu S, Pan Z, Deng J, Li X, Pan F (2019) Comparative Efficacy and Safety of Common Therapies in Keloids and Hypertrophic Scars: A Systematic Review and Meta-analysis. Aesthetic plastic surgery

Bayat A, McGrouther DA, Ferguson MWJ (2003) Skin scarring. BMJ: British Medical Journal 326: 88–92

Behrendt N (2004) The urokinase receptor (uPAR) and the uPAR-associated protein (uPARAP/Endo180): membrane proteins engaged in matrix turnover during tissue remodeling. Biological chemistry 385: 103–136

Bindea G, Mlecnik B, Hackl H, Charoentong P, Tosolini M, Kirilovsky A, Fridman WH, Pages F, Trajanoski Z, Galon J (2009) ClueGO: a Cytoscape plug-in to decipher functionally grouped gene ontology and pathway annotation networks. Bioinformatics (Oxford, England) 25: 1091–1093

Bock O, Schmid-Ott G, Malewski P, Mrowietz U (2006) Quality of life of patients with keloid and hypertrophic scarring. Archives of dermatological research 297: 433

Butler A, Hoffman P, Smibert P, Papalexi E, Satija R (2018) Integrating single-cell transcriptomic data across different conditions, technologies, and species. Nature Biotechnology 36: 411

Carney BC, Chen JH, Kent RA, Rummani M, Alkhalil A, Moffatt LT, Rosenthal DS, Shupp JW (2019) Reactive Oxygen Species Scavenging Potential Contributes to Hypertrophic Scar Formation. The Journal of surgical research 244: 312–323

Carthy JM (2018) TGFbeta signaling and the control of myofibroblast differentiation: Implications for chronic inflammatory disorders. Journal of cellular physiology 233: 98–106

Di Cera E (2009) Serine proteases. IUBMB life 61: 510–515

Elmansi AM, Awad ME, Eisa NH, Kondrikov D, Hussein KA, Aguilar-Pérez A, Herberg S, Periyasamy-Thandavan S, Fulzele S, Hamrick MW, McGee-Lawrence ME, Isales CM, Volkman BF, Hill WD (2019) What doesn’t kill you makes you stranger: Dipeptidyl peptidase-4 (CD26) proteolysis differentially modulates the activity of many peptide hormones and cytokines generating novel cryptic bioactive ligands. Pharmacology & therapeutics 198: 90–108

Fearmonti RM, Bond JE, Erdmann D, Levin LS, Pizzo SV, Levinson H (2011) The modified Patient and Observer Scar Assessment Scale: a novel approach to defining pathologic and nonpathologic scarring. Plastic and reconstructive surgery 127: 242–247

Ferguson MW, O’Kane S (2004) Scar-free healing: from embryonic mechanisms to adult therapeutic intervention. Philosophical transactions of the Royal Society of London Series B, Biological sciences 359: 839–850

Gschwandtner M, Mildner M, Mlitz V, Gruber F, Eckhart L, Werfel T, Gutzmer R, Elias PM, Tschachler E (2013) Histamine suppresses epidermal keratinocyte differentiation and impairs skin barrier function in a human skin model. Allergy 68: 37–47

Guo Y, Gupte M, Umbarkar P, Singh AP, Sui JY, Force T, Lal H (2017) Entanglement of GSK-3β, β-catenin and TGF-β1 signaling network to regulate myocardial fibrosis. Journal of molecular and cellular cardiology 110: 109–120

Gupta KK, Donahue DL (2017) Plasminogen Activator Inhibitor-1 Protects Mice Against Cardiac Fibrosis by Inhibiting Urokinase-type Plasminogen Activator-mediated Plasminogen Activation. 7: 365

Hammond TR, Dufort C, Dissing-Olesen L, Giera S, Young A, Wysoker A, Walker AJ, Gergits F, Segel M, Nemesh J, Marsh SE, Saunders A, Macosko E, Ginhoux F, Chen J, Franklin RJM, Piao X, McCarroll SA, Stevens B (2019) Single-Cell RNA Sequencing of Microglia throughout the Mouse Lifespan and in the Injured Brain Reveals Complex Cell-State Changes. Immunity 50: 253–271.e256

Hata A, Chen YG (2016) TGF-β Signaling from Receptors to Smads. Cold Spring Harbor perspectives in biology 8

Hinz B (2016) Myofibroblasts. Experimental eye research 142: 56–70

Hong SK, Choo EH, Ihm SH, Chang K, Seung KB (2017) Dipeptidyl peptidase 4 inhibitor attenuates obesity-induced myocardial fibrosis by inhibiting transforming growth factor-betal and Smad2/3 pathways in high-fat diet-induced obesity rat model. Metabolism: clinical and experimental 76: 4255

Hu MS, Longaker MT (2016) Dipeptidyl Peptidase-4, Wound Healing, Scarring, and Fibrosis. Plastic and reconstructive surgery 138: 1026–1031

Juillerat-Jeanneret L (2014) Dipeptidyl peptidase IV and its inhibitors: therapeutics for type 2 diabetes and what else? Journal of medicinal chemistry 57: 2197–2212

Kafka M, Collins V, Kamolz LP, Rappl T, Branski LK, Wurzer P (2017) Evidence of invasive and noninvasive treatment modalities for hypertrophic scars: A systematic review. Wound repair and regeneration: official publication of the Wound Healing Society [and] the European Tissue Repair Society 25: 139–144

Kaji K, Yoshiji H, Ikenaka Y, Noguchi R, Aihara Y, Douhara A, Moriya K, Kawaratani H, Shirai Y, Yoshii J, Yanase K, Kitade M, Namisaki T, Fukui H (2014) Dipeptidyl peptidase-4 inhibitor attenuates hepatic fibrosis via suppression of activated hepatic stellate cell in rats. Journal of gastroenterology 49: 481–491

Kanno Y (2019) The Role of Fibrinolytic Regulators in Vascular Dysfunction of Systemic Sclerosis. International journal of molecular sciences 20

Kant S, van den Kerckhove E, Colla C, van der Hulst R, Piatkowski de Grzymala A (2019) Duration of Scar Maturation: Retrospective Analyses of 361 Hypertrophic Scars Over 5 Years. Advances in skin & wound care 32: 26–34

Lal H, Ahmad F, Zhou J, Yu JE, Vagnozzi RJ, Guo Y, Yu D, Tsai EJ, Woodgett J, Gao E, Force T (2014) Cardiac fibroblast glycogen synthase kinase-3β regulates ventricular remodeling and dysfunction in ischemic heart. Circulation 130: 419–430

Lay AJ, Zhang HE, McCaughan GW, Gorrell MD (2019) Fibroblast activation protein in liver fibrosis. Frontiers in bioscience (Landmark edition) 24: 1–17

Leavitt T, Hu MS, Marshall CD, Barnes LA, Lorenz HP, Longaker MT (2016) Scarless wound healing: finding the right cells and signals. Cell and tissue research 365: 483–493

Lee HJ, Jang YJ (2018) Recent Understandings of Biology, Prophylaxis and Treatment Strategies for Hypertrophic Scars and Keloids. International journal of molecular sciences 1911

Lodyga M, Hinz B (2019) TGF-beta1 -A truly transforming growth factor in fibrosis and immunity. Seminars in cell & developmental biology

Lotia S, Montojo J, Dong Y, Bader GD, Pico AR (2013) Cytoscape app store. Bioinformatics (Oxford, England) 29: 1350–1351

Luecken MD, Theis FJ (2019) Current best practices in single-cell RNA-seq analysis: a tutorial. Molecular Systems Biology 15: e8746

Makrilakis K (2019) The Role of DPP-4 Inhibitors in the Treatment Algorithm of Type 2 Diabetes Mellitus: When to Select, What to Expect. International journal of environmental research andpublic health 16

Menou A, Duitman J, Crestani B (2018) The impaired proteases and anti-proteases balance in Idiopathic Pulmonary Fibrosis. Matrix biology: journal of the International Society for Matrix Biology 68-69: 382–403

Miao L, St Clair DK (2009) Regulation of superoxide dismutase genes: implications in disease. Free radical biology & medicine 47: 344–356

Muramatsu T (2010) Midkine, a heparin-binding cytokine with multiple roles in development, repair and diseases. Proceedings of the Japan Academy Series B, Physical and biological sciences 86: 410–425

Nabai L, Pourghadiri A, Ghahary A (2020) Hypertrophic Scarring: Current Knowledge of Predisposing Factors, Cellular and Molecular Mechanisms. Journal of Burn Care & Research 41: 48–56

Nishiguchi MA, Spencer CA, Leung DH, Leung TH (2018) Aging Suppresses Skin-Derived Circulating SDF1 to Promote Full-Thickness Tissue Regeneration. Cell reports 24: 3383–3392.e3385

Page MJ, Di Cera E (2008) Serine peptidases: classification, structure and function. Cellular and molecular life sciences: CMLS 65: 1220–1236

Qiu X, Hill A, Packer J, Lin D, Ma YA, Trapnell C (2017a) Single-cell mRNA quantification and differential analysis with Census. Nature methods 14: 309–315

Qiu X, Mao Q, Tang Y, Wang L, Chawla R, Pliner H, Trapnell C (2017b) Reversed graph embedding resolves complex single-cell developmental trajectories. bioRxiv: 110668

Rabieian R, Boshtam M, Zareei M, Kouhpayeh S, Masoudifar A, Mirzaei H (2018) Plasminogen Activator Inhibitor Type-1 as a Regulator of Fibrosis. 119: 17–27

Rawlings ND, Barrett AJ (1999) MEROPS: the peptidase database. Nucleic Acids Res 27: 325–331

Roberts AB, Sporn MB, Assoian RK, Smith JM, Roche NS, Wakefield LM, Heine UI, Liotta LA, Falanga V, Kehrl JH, et al. (1986) Transforming growth factor type beta: rapid induction of fibrosis and angiogenesis in vivo and stimulation of collagen formation in vitro. Proceedings of the National Academy of Sciences of the United States of America 83: 4167–4171

Rojahn TB, Vorstandlechner V, Krausgruber T, Bauer WM, Alkon N, Bangert C, Thaler FM, Sadeghyar F, Fortelny N, Gernedl V, Rindler K, Elbe-Bürger A, Bock C, Mildner M, Brunner PM (2020) Single-cell transcriptomics combined with interstitial fluid proteomics defines cell-type-specific immune regulation in atopic dermatitis. Journal of Allergy and Clinical Immunology

Schuliga M, Grainge C, Westall G, Knight D (2018) The fibrogenic actions of the coagulant and plasminogen activation systems in pulmonary fibrosis. Int J Biochem Cell Biol 97: 108–117

Schuliga M, Jaffar J, Harris T, Knight DA, Westall G, Stewart AG (2017) The fibrogenic actions of lung fibroblast-derived urokinase: a potential drug target in IPF. Scientific reports 7: 41770

Sen CK, Gordillo GM, Roy S, Kirsner R, Lambert L, Hunt TK, Gottrup F, Gurtner GC, Longaker MT (2009) Human skin wounds: a major and snowballing threat to public health and the economy. Wound repair and regeneration: official publication of the Wound Healing Society [and] the European Tissue Repair Society 17: 763–771

Shi S, Koya D, Kanasaki K (2016) Dipeptidyl peptidase-4 and kidney fibrosis in diabetes. Fibrogenesis & tissue repair 9: 1

Sidgwick GP, McGeorge D, Bayat A (2015) A comprehensive evidence-based review on the role of topicals and dressings in the management of skin scarring. Archives of dermatological research 307: 461–477

Snel B, Lehmann G, Bork P, Huynen MA (2000) STRING: a web-server to retrieve and display the repeatedly occurring neighbourhood of a gene. Nucleic Acids Res 28: 3442–3444

Stuart T, Butler A, Hoffman P, Hafemeister C, Papalexi E, Mauck WM, Stoeckius M, Smibert P, Satija R (2018) Comprehensive integration of single cell data. bioRxiv: 460147

Suzuki T, Tada Y, Gladson S, Nishimura R, Shimomura I, Karasawa S, Tatsumi K, West J (2017) Vildagliptin ameliorates pulmonary fibrosis in lipopolysaccharide-induced lung injury by inhibiting endothelial-to-mesenchymal transition. Respiratory research 18: 177

Szklarczyk D, Gable AL, Lyon D, Junge A, Wyder S, Huerta-Cepas J, Simonovic M, Doncheva NT, Morris JH, Bork P, Jensen LJ, Mering CV (2019) STRING v11: protein-protein association networks with increased coverage, supporting functional discovery in genome-wide experimental datasets. Nucleic Acids Res 47: D607–d613

Thielitz A, Vetter RW, Schultze B, Wrenger S, Simeoni L, Ansorge S, Neubert K, Faust J, Lindenlaub P, Gollnick HP, Reinhold D (2008) Inhibitors of dipeptidyl peptidase IV-like activity mediate antifibrotic effects in normal and keloid-derived skin fibroblasts. The Journal of investigative dermatology 128: 855–866

Trapnell C, Cacchiarelli D, Grimsby J, Pokharel P, Li S, Morse M, Lennon NJ, Livak KJ, Mikkelsen TS, Rinn JL (2014) The dynamics and regulators of cell fate decisions are revealed by pseudotemporal ordering of single cells. Nat Biotechnol 32: 381–386

Tredget EE, Shupp JW, Schneider JC (2017) Scar Management Following Burn Injury. Journal of burn care & research: official publication of the American Burn Association 38: 146–147

Uchida T, Oda T, Matsubara H, Watanabe A, Takechi H, Oshima N, Sakurai Y, Kumagai H (2017) Renoprotective effects of a dipeptidyl peptidase 4 inhibitor in a mouse model of progressive renal fibrosis. Renal failure 39: 340–349

Vallée A, Lecarpentier Y, Guillevin R, Vallée JN (2017) Interactions between TGF-β1, canonical WNT/β-catenin pathway and PPAR γ in radiation-induced fibrosis. Oncotarget 8: 90579–90604

Van Loey NE, Bremer M, Faber AW, Middelkoop E, Nieuwenhuis MK (2008) Itching following burns: epidemiology and predictors. The British journal of dermatology 158: 95–100

Vorstandlechner V, Laggner M, Kalinina P, Haslik W, Radtke C, Shaw L, Lichtenberger BM, Tschachler E, Ankersmit HJ, Mildner M (2020) Deciphering the functional heterogeneity of skin fibroblasts using single-cell RNA sequencing. FASEB journal: official publication of the Federation of American Societies for Experimental Biology 34: 3677–3692

Wang D, Zhang G, Chen X, Wei T, Liu C, Chen C, Gong Y, Wei Q (2018) Sitagliptin ameliorates diabetic nephropathy by blocking TGF-β1/Smad signaling pathway. International journal of molecular medicine 41: 2784–2792

