## Supplementary figures and images for "Single cell landscape of hypertrophic scars identifies serine proteases as key regulators of myofibroblast differentiation"

### Figure S1

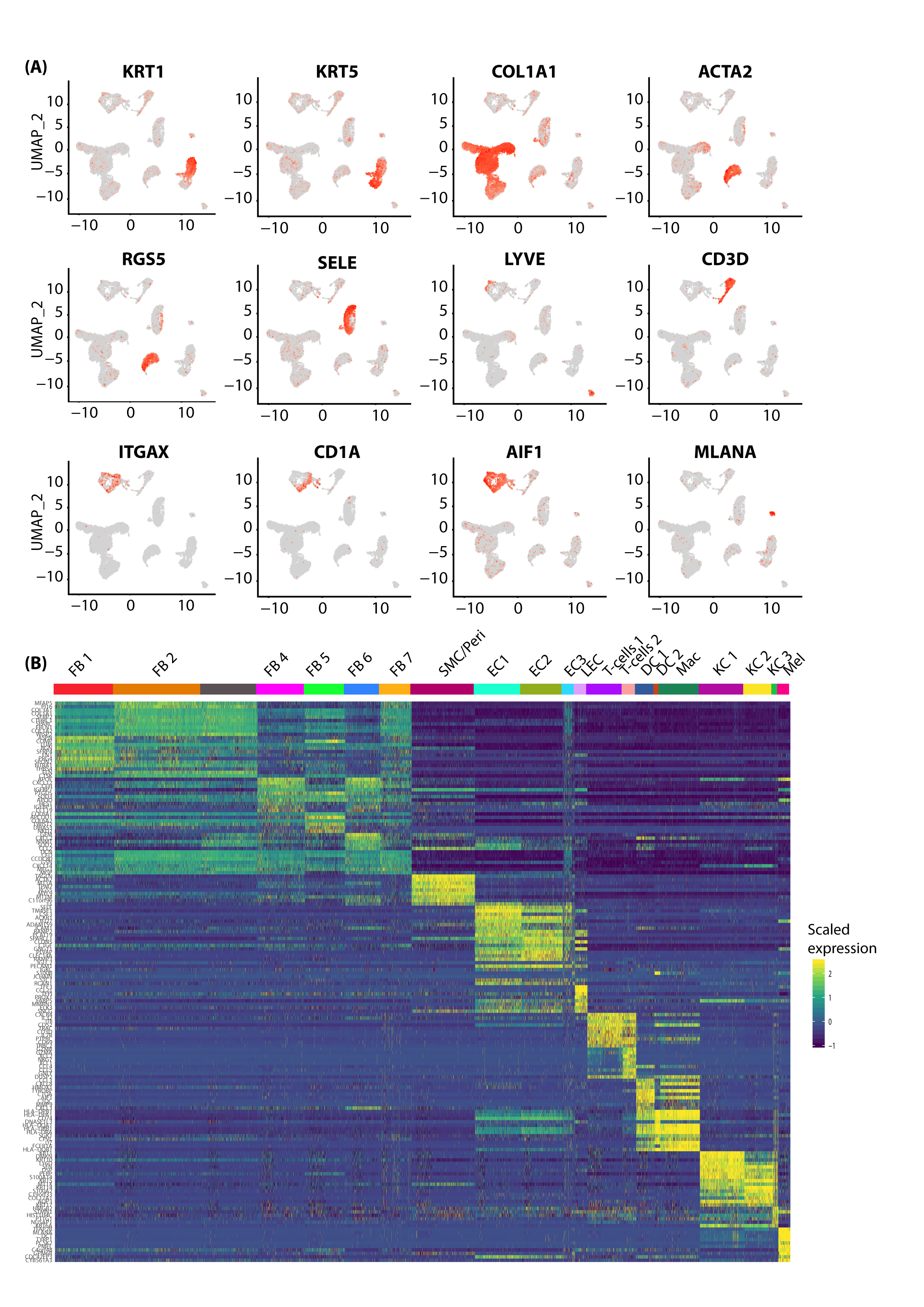

### Figure S2

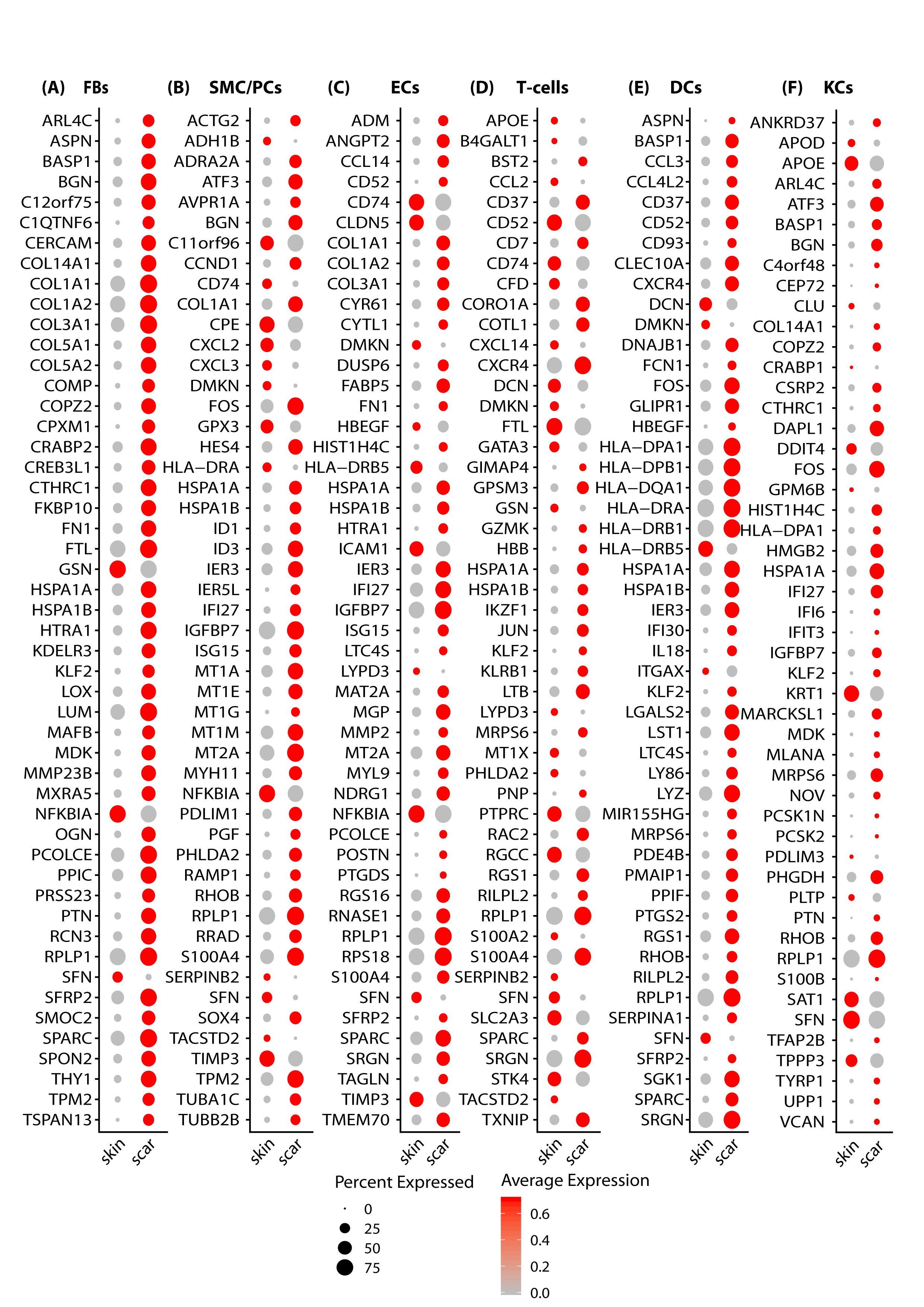

### Figure S3

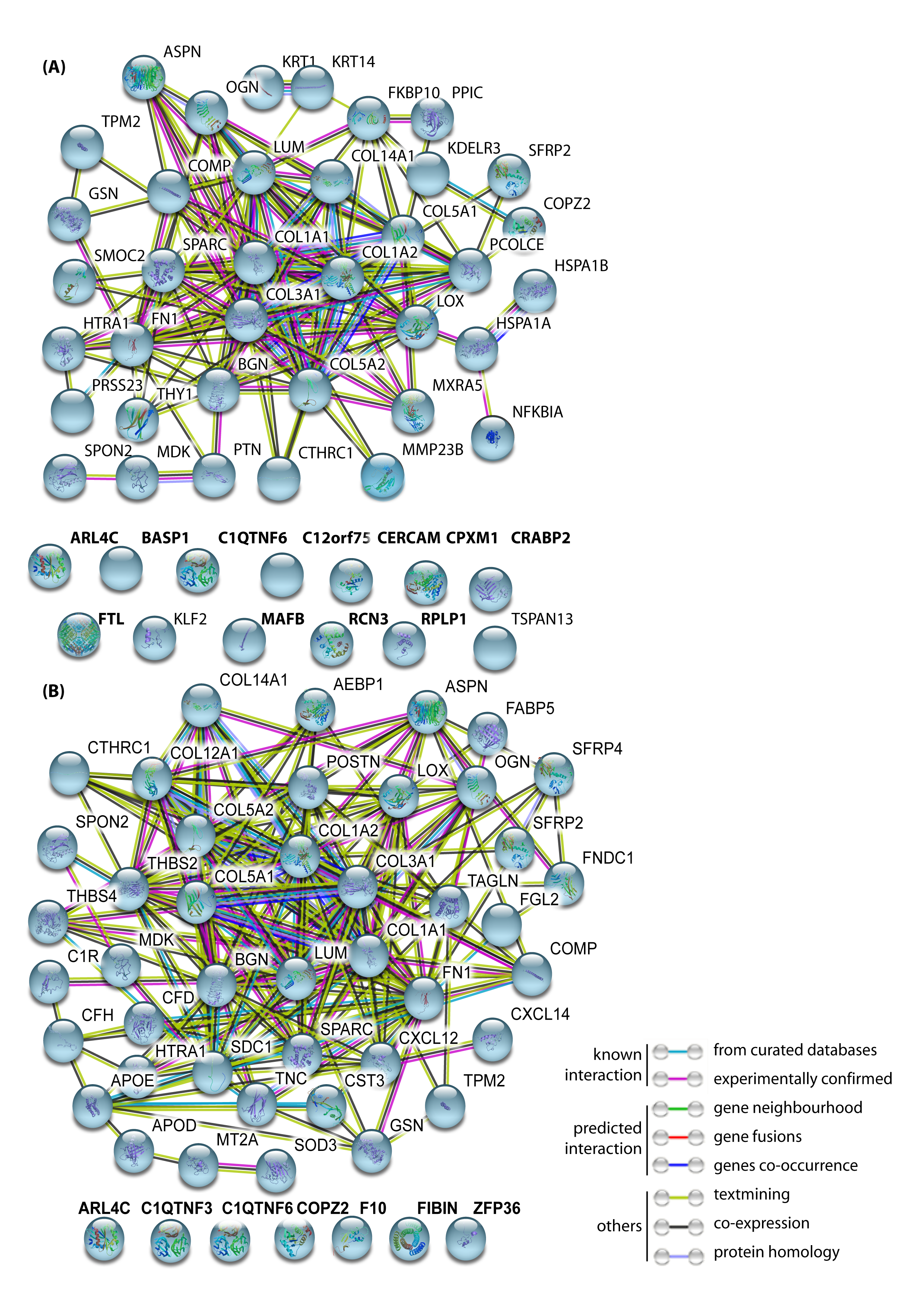

### Figure S4

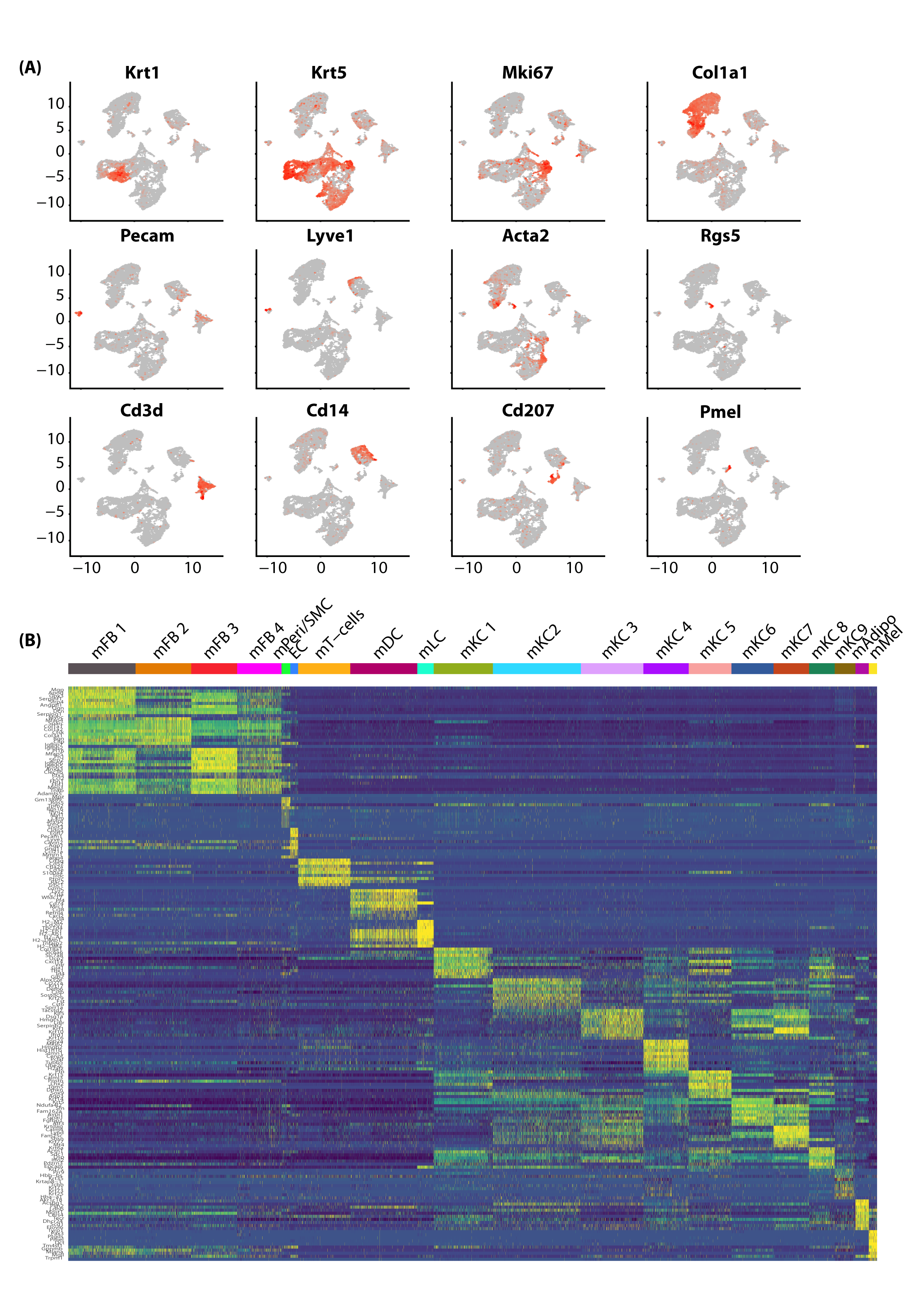

### Figure S5

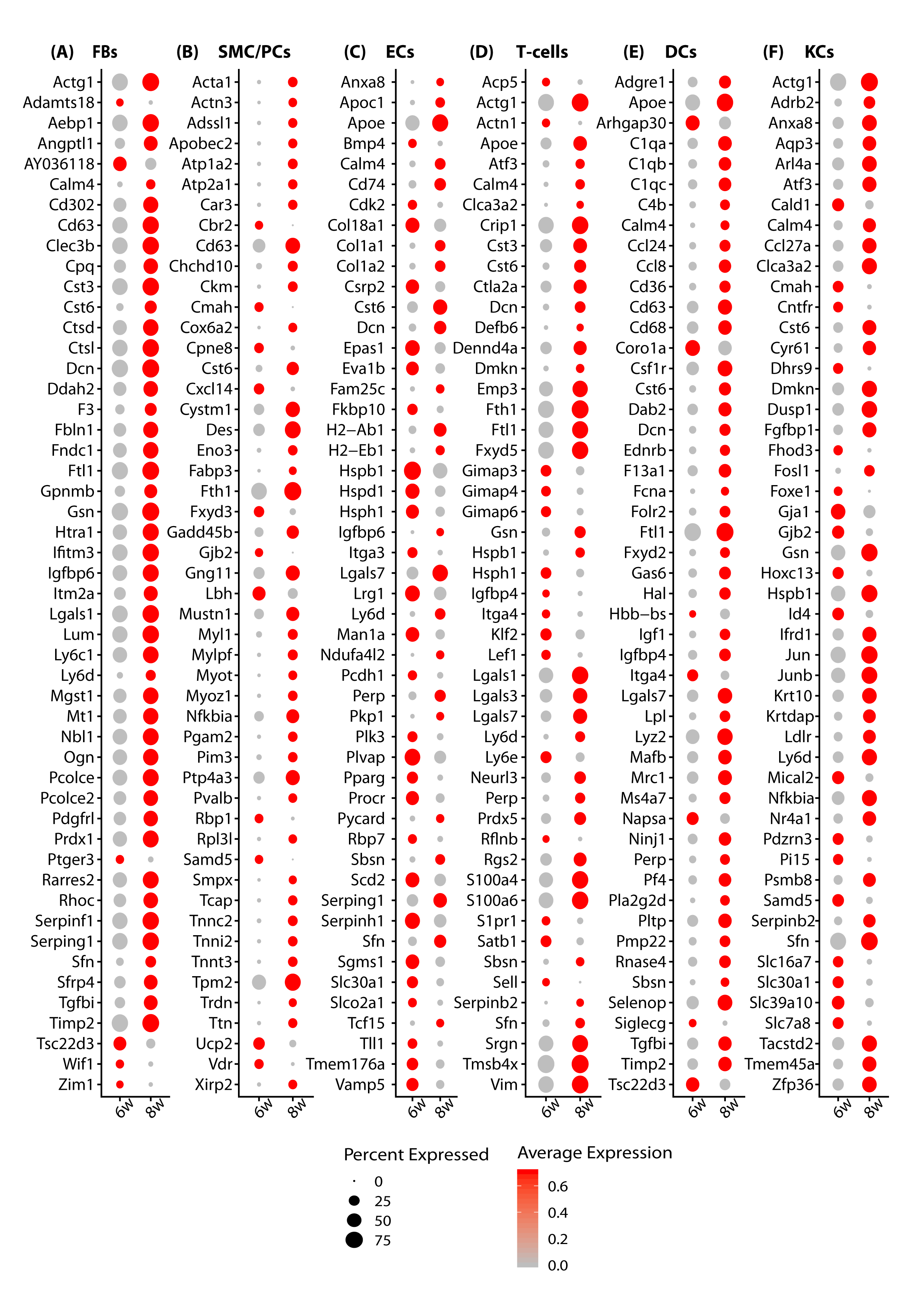

### Figure S6

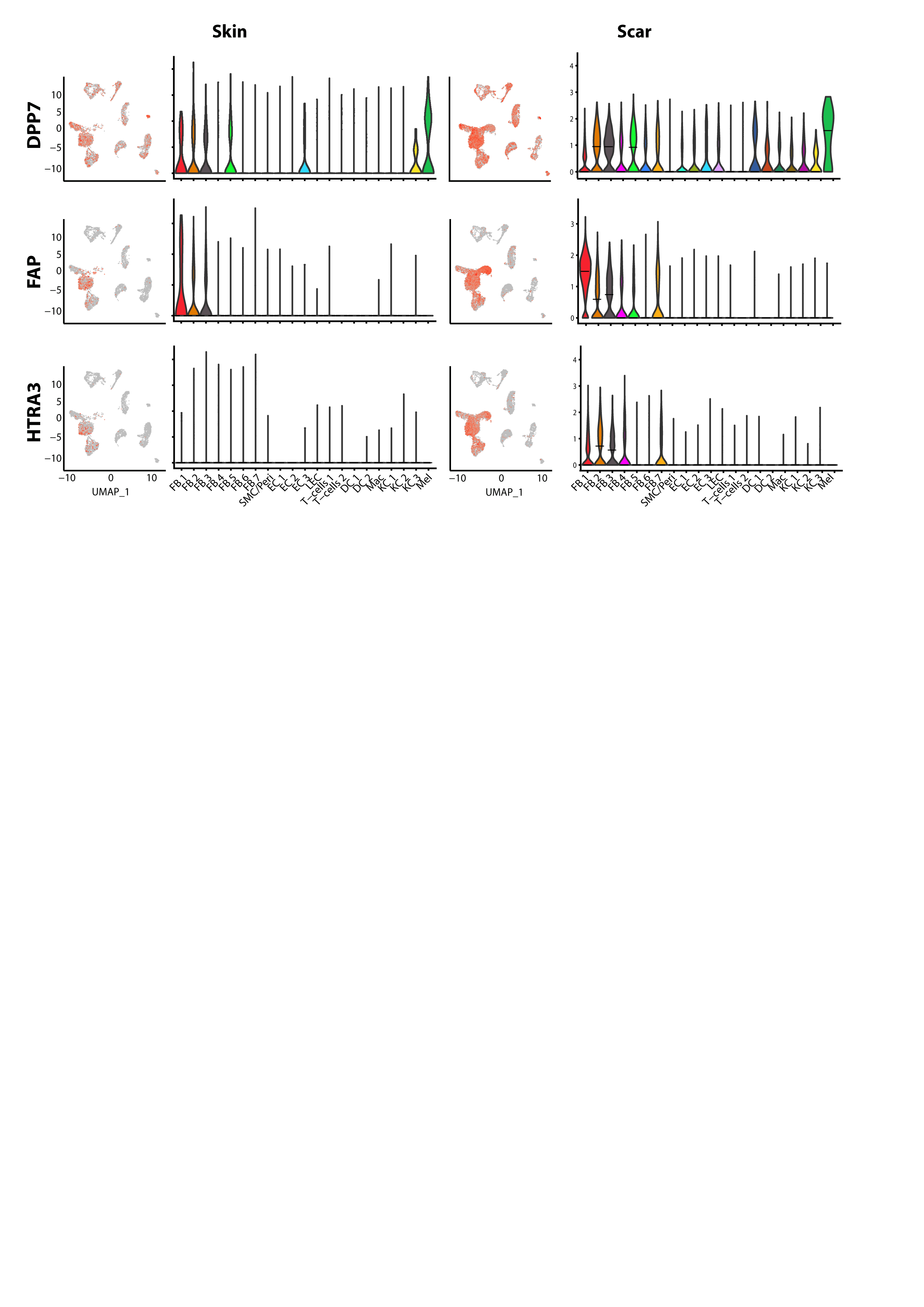

### Figure S7

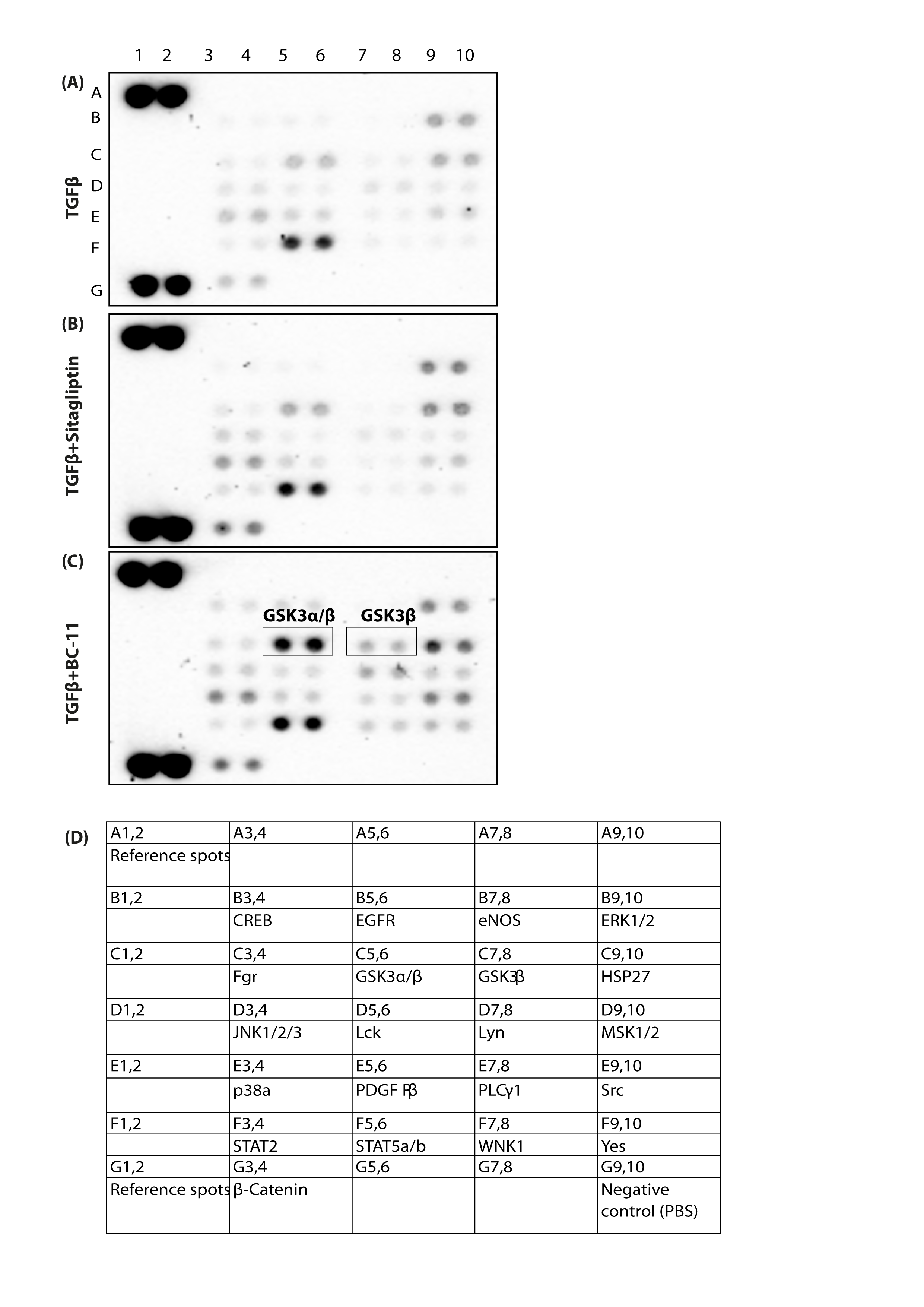

### Table S1

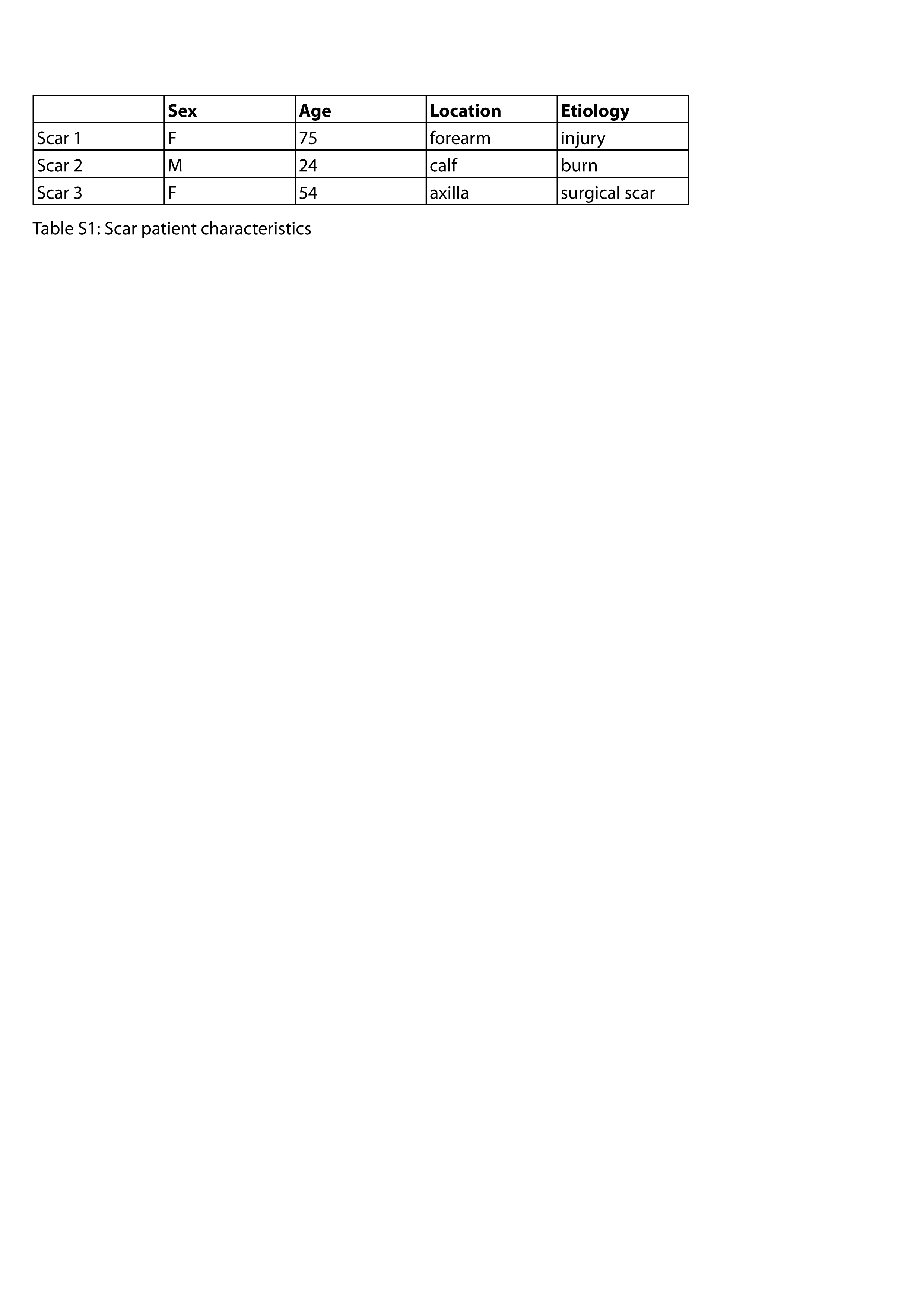

### Table S2

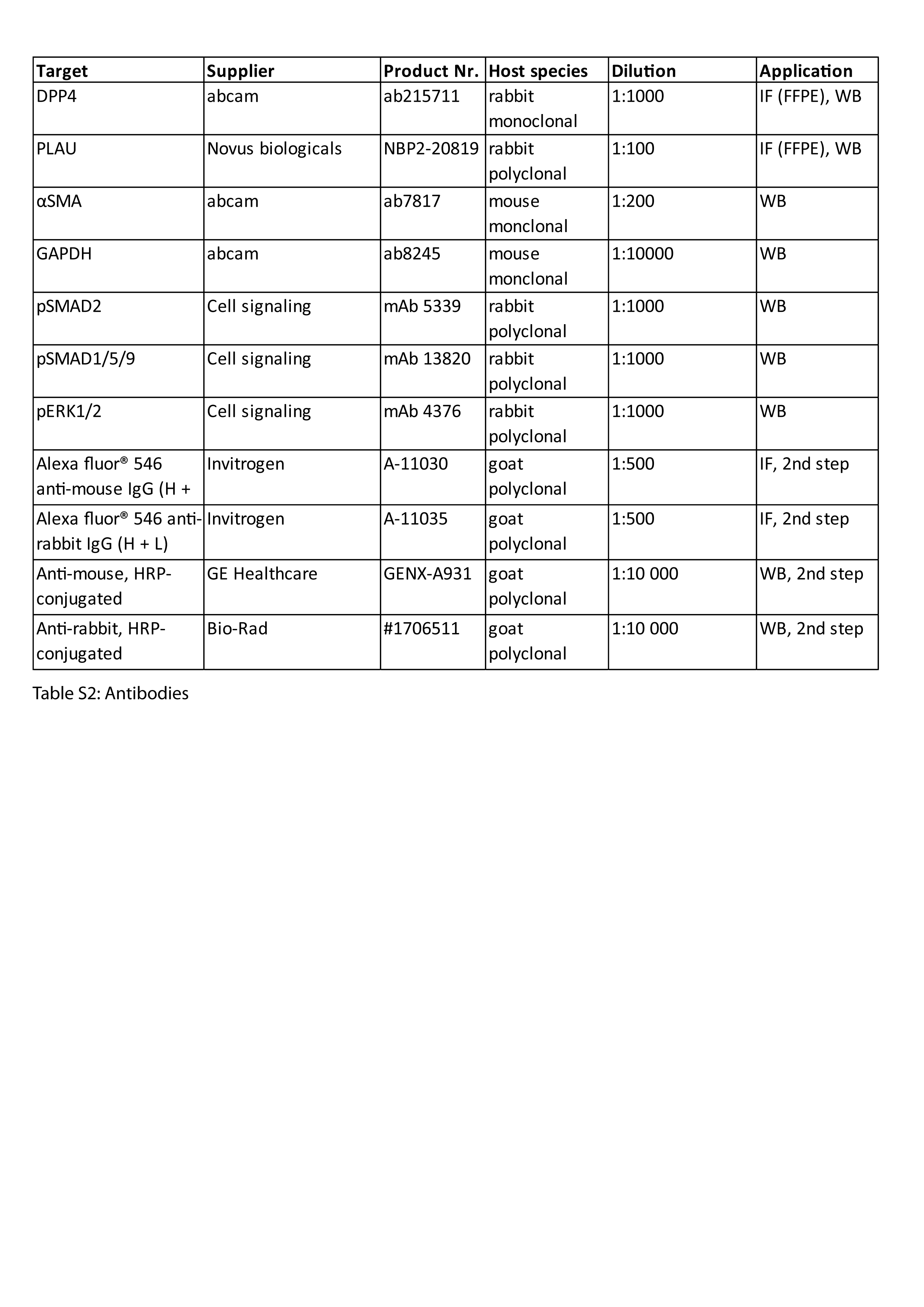
